# Genetic polymorphisms in a mate choice locus are maintained by balancing selection in a wild medaka population

**DOI:** 10.64898/2026.05.06.723183

**Authors:** Shingo Fujimoto, Taijun Myosho, Hirozumi Kobayashi, Hiroaki Aoyama, Iki Murase, Bayu K. A. Sumarto, Mitsuharu Yagi, Taiga Kunishima, Masatoshi Matsunami, Ryosuke Kimura

**Author notes:** Corresponding authors Shingo Fujimoto, Integrated Technology Center, University of the Ryukyus, Okinawa 9032720, Japan, (Present address) Ryosuke Kimura, Graduate School of Medicine, University of the Ryukyus, Okinawa 903-2720, Japan.

## Abstract

Sexual selection arises from individual differences in reproductive success, which can drive the maintenance of genetic polymorphisms in genes subject to balancing selection by the pleiotropic effects that trade-off between survival and reproduction. However, the extent to which sexual selection maintains genetic polymorphisms in wild populations remains unclear. Here, we explored on genomic signatures of balancing selection and selective sweep in the northern medaka, *Oryzias sakaizumii* in Japan by performing whole-genome resequencing of wild individuals. In addition, we re-evaluated the population genetic structure and admixture of *Oryzias latipes* and *O. sakaizumii* across the Japanese archipelago and detected genomic regions affected by introgression. Regions with signatures of selection from multiple statistics were located on eleven chromosomes. In particular, a region spanning 4.25 to 6.80 Mb on chromosome 18 showed high genetic diversity that could not be explained by sex differentiation or introgression from *O. latipes* in Eastern Japan. This pattern suggests that balancing selection maintains genetic polymorphisms in *O. sakaizumii*. Specifically, because a previously reported quantitative trait locus associated with female mating behavior overlaps with this region, we infer that sexual selection contributes to the maintenance of genetic polymorphism at this locus.

## Introduction

Genetic variation within a population provides information that can be used to assess and detect signatures of natural selection. When a favorable mutation undergoes positive selection, the fixation of the mutation leads to a reduction in genetic diversity via a hard selective sweep (Sabeti et al. 2002; Voight et al. 2006; Kimura et al. 2007; Ferrer-Admetlla et al. 2014). Conversely, balancing selection maintains genetic diversity through mechanisms such as heterozygote advantage, frequency-dependent selection (Hedrick 2007; Fijarczyk and Babik 2015), and antagonistic pleiotropy (Hedrick 1999; Fijarczyk and Babik 2015; Mérot et al. 2020; Abdul-Rahman et al. 2021). Genetic polymorphisms maintained by antagonistic pleiotropy have attracted increasing attention in mathematical models, evolutionary experiments, and population genomics, particularly with the development of nucleotide sequencing technologies (Wittmann et al. 2017; Zajitschek and Connallon 2018; Merot et al. 2020; Abdul-Rahman et al. 2021). These studies suggest that pleiotropic effects of genotypes on multiple fitness components can maintain genetic polymorphisms across broader regions of the genome than previously recognized (Wittmann et al. 2017; Zajitschek and Connallon 2018; Merot et al. 2020).

Sexual selection, which arises from individual differences in reproductive success, can promote balancing selection and thereby maintain genetic polymorphisms (Fijarczyk and Babik 2015; Wittmann et al. 2017; Zajitschek and Connallon 2018; Merot et al. 2020). Traits that evolve under sexual selection frequently incur antagonistic pleiotropic effects across various life-history components, including growth, survival, and reproduction (McBride et al. 2015; Simmons et al. 2017; Audzijonyte and Richards 2018; Mérot et al. 2020). Furthermore, phenotypic optimum often differs between males and females (Bonduriansky and Chenoweth 2009; Mank et al. 2017; Zajitschek and Connallon 2018). When the costs and benefits of traits depend on an individual’s ontogenetic stage, environmental conditions, and/or sex, the relative fitness of alleles at a locus can change even within a single population (Mank et al. 2017; Wittmann et al. 2017; Merot et al. 2020). Although numerous ecological studies have accumulated evidence of actual instances of sexual selection and selection pressures acting on phenotypes (Andersson and Simmons 2006; Wellenreuther et al. 2014; Radwan et al. 2016), the evolutionary consequences of sexual selection, particularly the maintenance of genetic polymorphisms at loci of sexual selection, remain poorly understood in wild populations.

To explore signature of selection in wild populations, the effect of demographic events such as population bottlenecks, population substructure, and admixture should be accounted for because the allele frequency spectrum across the genome is largely influenced by population demography (Excoffier et al. 2013; Ferrer-Admetlla et al. 2014; Fijarczyk and Babik 2015; Soni and Jensen 2024). Such demographic effects on allele frequency spectrum cause the false positive signal of selection. Consequently, when examining genetic diversity and selection signatures based on the empirical distribution of population genetic statistics, it is important to consider the influence of demographic events and to characterize the distribution of indicators of genetic diversity and selection.

In this study, we focused on two freshwater fish species in Japan, *Oryzias latipes* and *Oryzias sakaizumii*, collectively known as medaka. These two species have long served as model organisms in genetics, and geographic variation in secondary sexual traits has been documented, including fin length, male courtship frequency, female mate choice, and egg number (Kawajiri et al. 2014; Shinomiya et al. 2023; Fujimoto et al. 2024, 2026). In addition, the genetic architecture of interspecific variation has been evaluated using quantitative trait locus (QTL) analyses in interspecific F_2_ hybrids (Kawajiri et al. 2014, 2015; Yassumoto et al. 2020; Fujimoto et al. 2024). Molecular basis of male specific morphology and mating behavior in both sexes has also examined by gene knock-out strains (e.g., Ogino et al. 2023, Nishiike et al. 2021). The information on genetic architecture associated with sexual dimorphism, mating behaviors, and reproductive traits provides a foundation for identifying chromosomal regions potentially shaped by sexual selection.

Despite these advances, few studies have identified genes under selection in wild medaka populations. Previous selection scan in this fish has focused only on a single collection site of *O. latipes* with small sample sizes (Spivakov et al. 2014), and genome-wide evidence of balancing selection remains absent. Additionally, although the possibility of adaptation to high-latitude environments in the northern medaka *O. sakaizumii* has been hypothesized (Shinomiya et al. 2023; Fujimoto et al. 2024, 2026), selection scans for this species have not yet been conducted. Therefore, we investigated the signature of selection specifically in wild *O. sakaizumii* population.

Wild medaka populations in Japan have experienced secondary contact and admixture in their natural habitats (Takehana et al. 2016; Katsumura et al. 2019). Recent studies suggest that admixture between the two *Oryzias* species occurred across a geographically broader range than previously recognized (Fujimoto et al. 2022). Hence, we re-evaluated the population genetic structure and admixture across the Japanese archipelago using whole-genome resequencing. The analysis of selection signature and introgressed regions allows that we identify regions with high genetic diversity due to balancing selection rather than introgression. Based on the detected selection signatures, we discuss the possibility that sexual selection maintains genetic polymorphisms at the previously reported QTLs associated with female mating behavior. We also discuss how male-female behavioral interactions during the fertilization may contribute to the maintenance of genetic polymorphisms.

## Materials and Methods

### Fish populations and field collection

Medaka, or the *Oryzias latipes* species complex, comprising *O. latipes*, *O. sinensis*, and *O. sakaizumii*, are small freshwater fish distributed in low-lying wetlands throughout Japan, Korea, and China (Iwamatsu 2006, Fig. 1a). Population genetic analyses using nuclear SNVs have shown that the Japanese populations of the *O. latipes* species complex consist of several genetically distinct groups (Katsumura et al. 2019, Fujimoto et al. 2022). *Oryzias latipes*, which is distributed along the coasts of the Pacific and the East China Sea, exhibits a roughly southwest-to-northeast genetic cline (Katsumura et al. 2019, Fujimoto et al. 2022). The northern medaka, *O. sakaizumii*, is mainly distributed along the coast of the Sea of Japan (Fig. 1a). This genetic group of the *O. latipes* species complex has been used as a model organism in genetics. Reference genome sequences have been constructed for four genetically divergent strains (Ichikawa et al. 2017, Dong et al. 2024).

**Figure 1.**
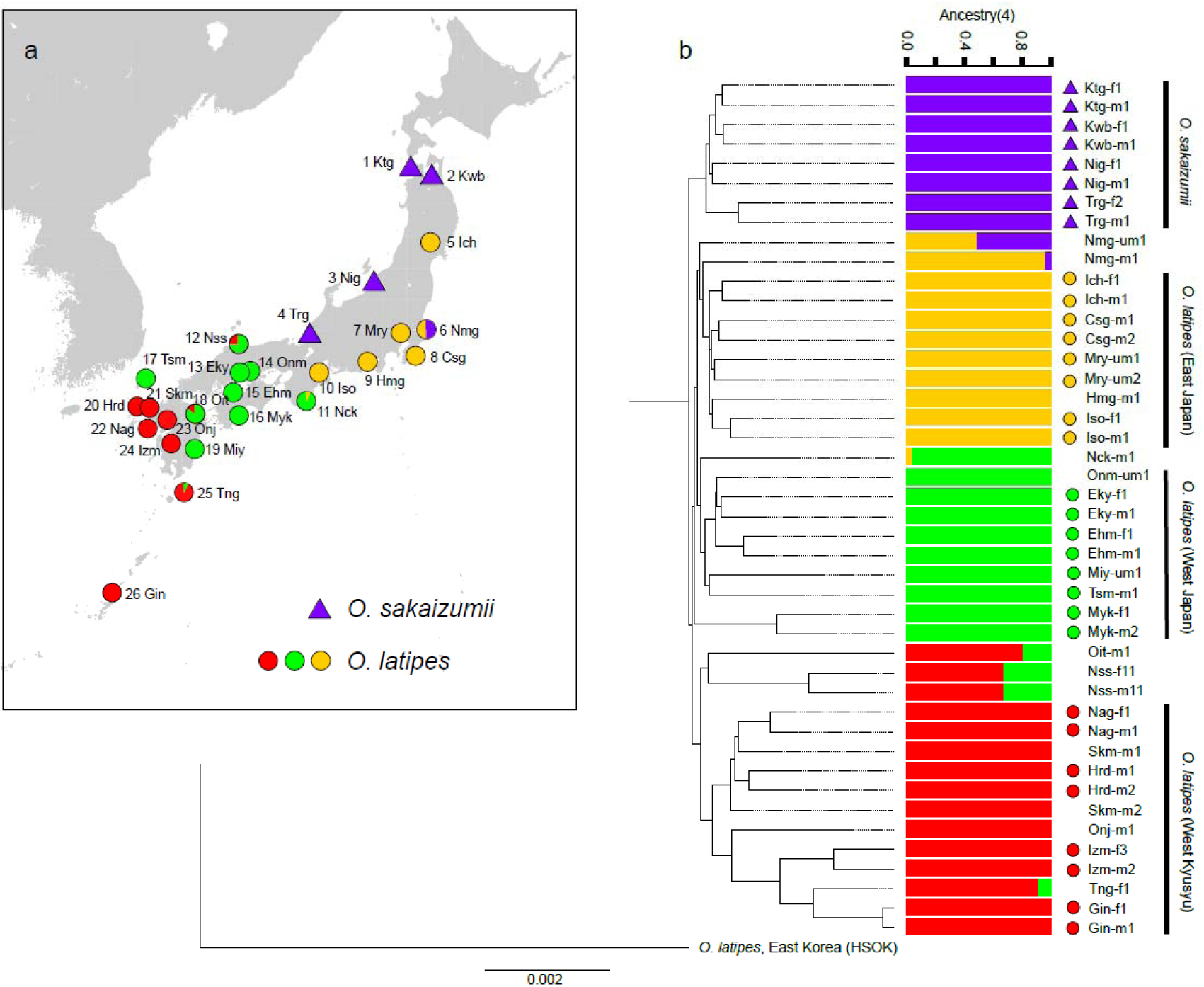
(a) Geographic locations of collection sites. Solid circles and triangles on the map represent *O. sakaizumii* and *O. latipes*, respectively. Colors correspond to the clusters in the ADMIXTURE analysis. (b) Neighbor-joining tree based on the total whole-genome sequences and results of ADMIXTURE analysis for *K* = 4. Individuals marked with a circle or triangle were used in the admixture analysis represented in Figure 2. For the abbreviations of the wild populations, see Tables S1 and S2.

To analyze the population structure of wild medaka populations, we collected wild individuals throughout the distribution of the *Oryzias latipes* species complex in Japan (Fig. 1a, Table S1). The collection sites covered the major habitats of medaka in Japan. At most sites, we obtained whole-genome sequence data from more than two individuals per site (Tables S1, S2), including four sites for *O. sakaizumii* and 22 sites for *O. latipes*. Additionally, to examine genetic diversity within *O. sakaizumii*, we obtained sequence data from 8 or 16 individuals per site (*N* = 40). We also used the publicly available sequences of the inbred strain, HSOK (Spivakov et al. 2014, DRA000588), which originated in Korea and was used as an outgroup for the analysis of the Japanese populations.

### Library construction, sequencing, and genotyping

For these wild-collected individuals, live fish were anesthetized with MS-222 (tricaine methanesulfonate, FUJIFILM Wako Pure Chemical Corporation, Osaka, Japan) and then fixed in 99% ethanol. Genomic DNA was extracted from ethanol-fixed pectoral fin samples using a DNeasy Kit (Qiagen Inc., Hilden, Germany) according to the manufacturer’s instructions. Sample libraries for whole-genome re-sequencing were prepared using a TruSeq DNA PCR-free Library Prep Kit (Illumina Inc., San Diego, CA, USA) according to the manufacturer’s instructions. The DNA concentration of each library was measured using a StepOnePlus real-time PCR system (Thermo Fisher Scientific, Waltham, MA, USA) with a KAPA Library Quantification Kit (Kapa Biosystems, Inc., Woburn, MA, USA). Five or six individual libraries were pooled in equal amounts and sequenced using a HiSeq X system (Illumina, PE, 150 bp). Sequencing was performed by Macrogen Japan (Tokyo, Japan). Sequences were deposited in the DDBJ Sequence Read Archive (DRA) under the accession number PRJDB20255.

For each sample, the Illumina adapter sequences were removed, and quality filtering was performed using fastp ver. 0.20.0 with the options “--detect_adapter_for_pe, --cut_front”. The remaining reads were aligned to the medaka HNI strain reference genome (GCA_002234675.1; ASM223467) using the Burrows-Wheeler Alignment tool, BWA mem ver. 0.7.17 (Li, 2013). HNI was inbred strain, originally collected from the *O. sakaizumii* wild population in Niigata prefecture, and maintained by NBRP Medaka (https://shigen.nig.ac.jp/medaka/). After mapping, the output files were converted to Binary Alignment/Map (BAM) format using SAMtools ver. 1.7 (Danecek et al. 2021).

SNVs and InDels were called following the best practice guidelines of the Genome Analysis Tool Kit (GATK ver. 3.8.0) (Auwera and O’Connor 2020). We then filtered out variants based on the following criteria: “QD < 2.0 || FS > 60.0 || MQ < 40.0 || MQRankSum < −12.5 || ReadPosRankSum < −8.0” for SNVs, and “QD < 2.0 || FS > 200.0 || ReadPosRankSum < −20.0” for InDels. Based on the filtered SNVs and InDels, we recalibrated the mapping quality of each read and extracted sequences including variant and non-variant sites in vcf format using the “GenotypeGVCFs” command in GATK with the “--includeNonVariantSites” option selected.

SNV data were prepared using bcftools (Danecek et al. 2021). For each individual, sequence positions with depths (DP) between 10 and 200, mapping quality (MQ) ≥ 40, and quality by depth (QD) ≥ 2.0 were included in the analysis. Individual vcf files were merged into a multi-sample vcf file, and sequence positions that were genotyped in all individuals were extracted using the “bcftools view” command with the “NO_MISSING=0” option. Sites containing InDels were then excluded.

### Analysis of population structure and admixture

In total, 271085758 bases at non-variant and variant positions were sequenced from 45 individuals, including eight *O. sakaizumii*, 36 *O. latipes* in Japan, and one inbred strain of *O. latipes* from eastern Korea (HSOK) (Table S2). Phylogenetic relationships were estimated by the neighbor-joining method using a genetic distance matrix calculated using VCF2Dis ver. 1.47 (Xu, 2025, *Gigascience*, in press). The distance matrix was converted to the nexus format using the “writeDist” function of the “phangorn” package ver. 2.12.1 (Schliep 2011). The phylogenetic tree and network were constructed using SplitsTree ver. 4.15.1 (Huson and Bryant 2006) and Figtree ver. 1.4.3 (https://github.com/rambaut/figtree/). The branch between the Korean HSOK *O. latipes* from Korea and Japanese populations of *O. sakaizumii* and *O. latipes* was used as the root. To estimate population structure, we used ADMIXTURE ver. 1.3.0 (Alexander et al. 2009). ADMIXTURE was run with the number of clusters (*K*) ranging from 1 to 10, and statistical support for the different numbers of clusters was evaluated based on the lowest cross-validation error.

To estimate past admixture between populations, *Patterson’s D* were calculated for a dataset of 33 individuals, including eight individuals from each population (*O. sakaizumii*: Osak; *O. latipes*: Western Kyushu (WK), Western Japan (WJ), and Eastern Japan (EJ)) and one HSOK individual as an outgroup. In this analysis, individuals with multiple ancestries in the ADMIXTURE results, presumably due to recent migration or admixture, were excluded.

To further investigate population admixture, we applied the *F*-statistics framework (Durand et al. 2011, Patterson et al. 2012, Peter 2016). We treated HSOK as an outgroup and performed an *ABBA-BABA* test to calculate *Patterson’s D* statistic using “qpDstat” ver. 980 in ADMIXTOOLS (Patterson et al. 2012), which estimates admixture between four populations. For this analysis, SNVs that were polymorphic across the four populations were extracted using the “admixr” package (Petr et al. 2019). We reconstructed an admixture graph under an assumed demographic scenario using the “qpGraph” command in ADMIXTOOLS2 (Maier et al. 2023). We selected the best-fitting model for which the *F_4_* statistic for all the population combinations was less than three. In addition, introgressed loci were identified using *F_d_* statistics (Martin 2015), a modified version of *Patterson’s D* designed for small genomic regions. We calculated *F_d_* statistics using the “Dinvestigate” command in Dsuite ver. 0.5 r46 (Malinsky et al. 2021) in non-overlapping windows of 50 SNVs. Introgressed regions from the EJ population to the *O. sakaizumii* population were defined as windows the upper 0.5% tail of the genome-wide *F_d_*distribution.

### Estimation of divergence generations

Divergence between populations was estimated using MSMC2 v.2.0.0 (Schiffels et al. 2020). To create a mask file specifying the genomic regions to be analyzed in the HNI reference genome sequence, we used Snpable with the option “35mer, *r* = 0.5” (Heng Li, https://lh3lh3.users.sourceforge.net/snpable.shtml). Focusing on sites where two individuals were sequenced from the same site (Fig. 1b), we calculated coalescence rates using four individuals (i.e., eight haplotypes) from two populations or from two *O. sakaizumii* collection sites. A VCF file containing four diploid individuals was converted using the script generate_multihetsep.py in msmc-tools (Stephan Schiffels, https://github.com/stschiff/msmc-tools/tree/master), and coalescence rates were calculated using the default settings. The relative cross-population coalescence rate (rCCR) was then calculated, and the time point at which the CCR exceeded 0.5 was used as an indicator of population divergence (Schiffels and Durbin 2014). Coalescence rates were converted to divergence times in generations, assuming a mutation rate per generation of 3.50 × 10^−9^ per generation (Sutra et al. 2019, Malinsky et al. 2018) and a generation time of 1 year.

### Genomic scan to detect selection signatures

Focusing on biallelic SNVs in *O. sakaizumii*, π, *Tajima’s D* (Tajima 1989), and genetic differentiation between males and females (intersexual *FST*) were calculated using pixy (Korunes and Samuk 2020) and VCF-kit (Cook 2017). Each chromosome was divided into non-overlapping 50,000 bp windows. Windows with fewer than 10,000 bases sequenced in total across individuals were excluded from subsequent genomic scan analyses. We identified the top (or bottom) 0.5% tails of the genome-wide distributions for the statistics (Fig. S1). Then, (1) windows with high π and high *Tajima’s D*, (2) windows with low π and low *Tajima’s D*, and (3) windows with high *Tajima’s D* and high intersexual *FST* were extracted (Fig. S1a-c).

We also calculated the integrated haplotype score (iHS) in *O. sakaizumii*, which is based on the allele frequency and linkage disequilibrium between SNPs. Individual haplotypes were estimated from 40 *O. sakaizumii* and 42 *O. latipes* individuals (Table S2) using the “phase_common_static” command in Shapeit4 (Delaneau et al. 2019). Biallelic SNPs with a minor allele frequency greater than 5% in the 40 *O. sakaizumii* individuals were extracted and Z-transformed iHS for each SNP were calculated using Selscan ver. 2.0.0 (Szpiech 2024). For the iHS calculation, we used physical position information instead of fine-scale recombination map, which is not available for medaka. The top (positive) 0.25% and bottom (negative) 0.25% of Z-transformed iHS values in the genome-wide distribution were treated as outliers (Fig. S1d).

### Gene ontology analysis and phenotype association

To explore the molecular functions of genes located within the regions identified in the genomic scan, we conducted GO enrichment analysis. Gene names and GO annotations were downloaded from Ensembl release 109 (https://feb2023.archive.ensembl.org/) for the *O. latipes* HdrR genome (ASM223467v1, Ichikawa et al. 2017). The HdrR strain was inbred strain of orange mutant, which maintained in National BioResource Project (NBRP Medaka: https://shigen.nig.ac.jp/medaka/). The genetic composition of nuclear genome in the HdrR strain similar to that of the Western Japan population of *O. latipes* (WJ) and the Eastern Japan population of *O. latipes* (EJ) (Fujimoto et al. 2022). Since the HdrR genome is better annotated than the HNI genome of *O. sakaizumii* (ASM223471v1, Ichikawa et al. 2017), we used the HdrR genome for this analysis. For example, the HNI reference lacks a comprehensive ortholog database, and gene symbols are undetermined for many Ensembl gene IDs, whereas these annotations are available for the HdrR strain. A list of orthologs between the HNI and HdrR reference sequences was generated using OrthoFinder (Emms and Kelly 2019) and a custom R script (Table S3). GO enrichment analysis was then performed using HdrR gene annotations and ShinyGO ver. 0.82 (Ge et al. 2020). Enrichment analyses were performed separately for Ensembl gene IDs located in regions with signatures of introgression and selection. Statistical significance was assessed at *FDR* < 0.01.

Detection of selection signatures, such as *Tajima’s D* and iHS, depends on haplotype structures in the wild population. The population genetic statistics tended to be inflated in regions with low recombination rates (Stevison and McGaugh 2020), such as the near centromere regions (Peñalba and Wolf 2020). Hence, we inferred the centromere region in HNI reference sequence (ASM223471v1), we used the synteny block information of the chromosomal regions between the HNI and Hd-rR (Ansai et al. 2023, Table S4). Physical position of synteny block between HNI and Hd-rR was estimated using ntSynt, v.1.0.0 (Coombe et al. 2024). We examined the extent and overlap of the 1 Mb region near the centromere region (Table S4).

We also examined the overlap between genomic scan windows and phenotype associations, based on previous QTL analyses (e.g., Kawajiri et al. 2014, Fujimoto et al. 2024). The original genetic markers used in the QTL study were mapped to the *O. latipes* HdrR reference genome (Fujimoto et al. 2024). Therefore, genomic positions of the genetic markers were lifted from the HdrR reference (ASM223467v1) and converted to the updated reference genome (ASM223471v1) using the LiftOver software package (Hinrichs et al. 2006). This liftover was performed with “flo” (https://github.com/wurmlab/flo/, Pracana et al. 2017), a pipeline incorporating UCSC tools and BLAT (Kent et al. 2002).

## Results

### Population structure and past admixture

Phylogenetic relationships were inferred using the neighbor-joining method based on whole-genome sequence data from 45 individuals, including *O. latipes* and *O. sakaizumii* collected from 26 sites across Japan as well as the inbred HSOK strain originating from Korea. The Japanese populations were genetically separated into two species, *O. latipes* and *O. sakaizumii* (Fig. 1, Tables S1, S2). ADMIXTURE analysis supported the presence of four genetic populations (Fig. 1b, *K* = 4, CV error = 0.40, Fig. S2, Table S5), which correspond to the clusters in the neighbor-joining tree. Individuals of *O. sakaizumii* collected from four sites were grouped into a single population (hereafter referred to as Osak), while *O. latipes* was separated into three populations: Western Kyushu (WK), Western Japan (WJ), and Eastern Japan (EJ). Some individuals of *O. latipes* collected from Nmg (Namegata), Nck (Nachikatsuura), Nss (Nishihamasata), Oit (Oita), and Tng (Nakatane, Tanegashima) showed multiple genetic ancestries in the ADMIXTURE analysis (Fig. 1, Fig. S2), likely reflecting recent migration. After removing these admixed individuals, we confirmed that the clusters in the neighbor-joining tree did not change (Fig. 1, Fig. S3). In the subsequent admixture analysis, we used only individuals showing a single ancestry, which are marked with circles or triangles in Fig. 1b.

Genetic distances between populations were evaluated using outgroup *F_3_* statistics with the eastern Korean population, HSOK, as the outgroup. The closest populations were WJ and EJ, and the most distant were WK and Osak (Fig. 2a). Based on MSMC2, the average population divergence times were 2.97 × 10^6^ ± 0.57 × 10^6^ (SD) generations between WK and Osak, 1.01 × 10^6^ ± 0.40 × 10^6^ between WJ and WK, 1.15 × 10^6^ ± 0.50 × 10^6^ between EJ and WK, and 0.48 × 10^6^ ± 0.16 × 10^6^ between EJ and WJ (Fig. S4). The average divergence time among the four *O. sakaizumii* collection sites (Tsuruga, Niigata, Aomori, and Kitatsugaru) was estimated to be 3.06 × 10^4^ ± 3.17 × 10^4^ generations. The estimated number of generations within *O. sakaizumii* was two orders of magnitude smaller than those between *O. sakaizumii* and *O. latipes* (Fig. S4c, d).

**Figure 2.**
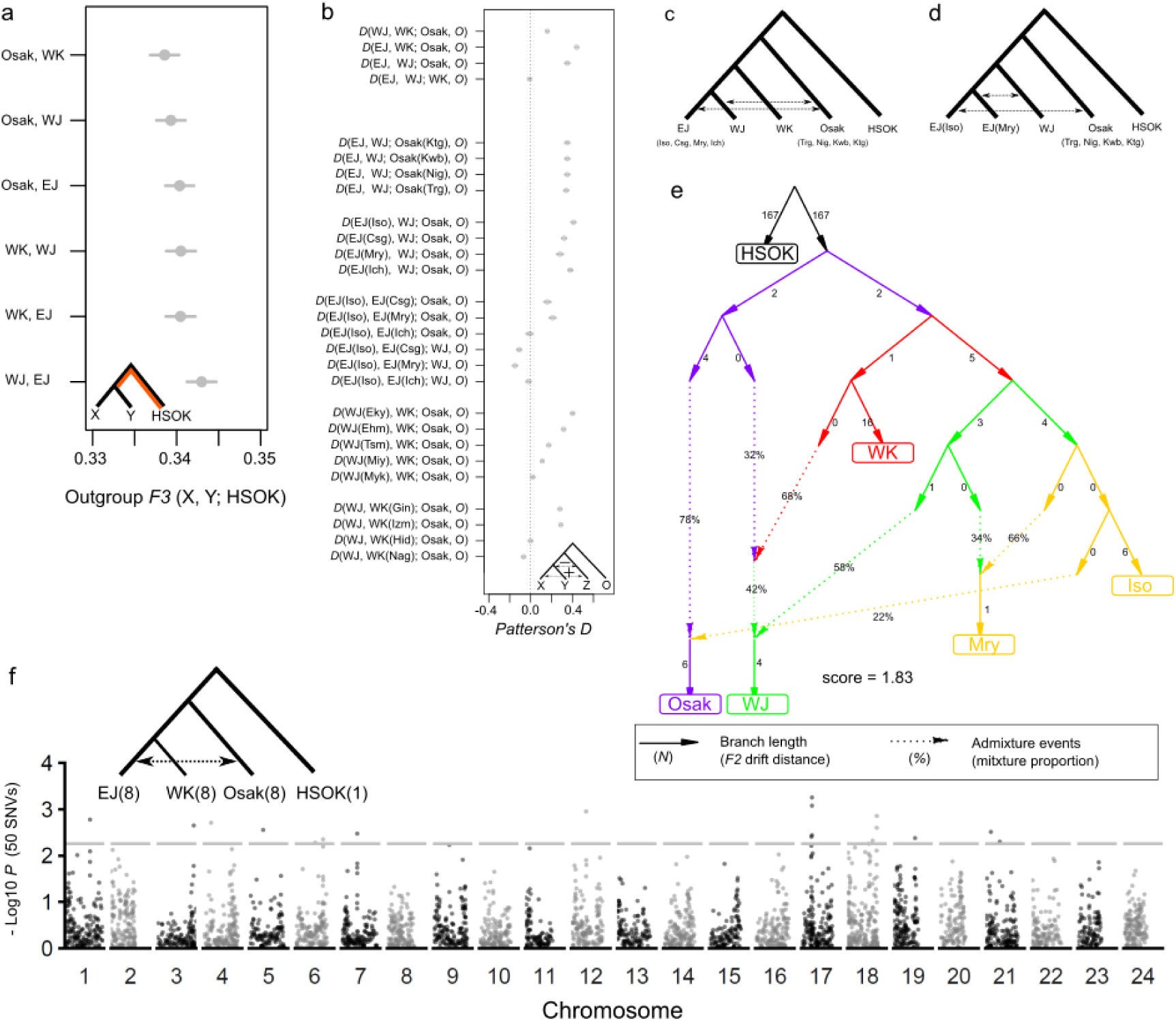
Admixture analyses focusing on *O. sakaizumii* and the Eastern Japan population of *O. latipes*. (a) Outgroup *F_3_* statistics identifying the most closely related populations. (b) Results of *Patterson’s D* across all possible combinations among five populations. (c, d) Schematic representations of unadmixed and admixed populations based on *Patterson’s D*. (e) Admixture graph best fitting the data. Branch lengths (*F_2_* drift distances) are shown as solid lines, and admixture events are indicated by dotted lines with mixture proportions. (f) Introgressed regions examined by a genomic scan of *F_d_* statistics in 50-SNV windows (Martin 2015) in the admixture analyses dataset. Y axis represented the −log10 transformed *P* value of empirical distribution in *F_d_*. Positive values indicated the introgression between EJ and Osak. The numbers in parenthesis showed the total sequenced individuals of each population.

To test for past admixture, we performed the *ABBA-BABA* test (Patterson’s *D* statistic). We found that *D* (EJ, X; Osak, HSOK), where X is WJ or WK, was significantly positive (|Z| = 21.50-30.97), indicating admixture between Osak and EJ (Fig. 2b–d, Table S6). When *D* (EJ, WJ; Osak, HSOK) was calculated by treating Osak separately for each collection site, the *D* values were constant. In contrast, when EJ was treated separately by collection site, the *D* values varied. Specifically, *D* (EJ(Iso), EJ2; Osak, HSOK) and *D* (EJ(Iso), EJ2; WJ, HSOK), where EJ2 was EJ(Csg) or EJ(Mry), showed positive and negative values, respectively. These results indicate that EJ(Iso)(and EJ(Ich)) interacted with Osak, whereas EJ(Csg) and EJ(Mry) interacted with WJ. The comparison between WJ and WK, i.e., *D* (WJ, WK, Osak HSOK), also yielded a positive value. Together, these patterns suggest the presence of local admixture between populations, varying by collection site.

We further reconstructed the history of the admixture events using qpGraph. In this analysis, we ultimately included five populations (i.e., HSOK, Osak, WK, WJ, EJ(Iso), and EJ(Mry)) in the model, because the EJ subpopulations appeared to interact differently with other populations. The resulting graph had a score of 1.83 and the maximum *F* combination was |Z| = 1.18 (HSOK, Osak; HSOK, Iso) (Fig. 2e, Fig. S5). The graph indicated that the interaction between Osak and EJ(Iso) occurred in the direction from EJ(Iso) to Osak, and that EJ(Mry) had received gene flow from the ancestral lineage of WJ. WJ appears to have experienced gene flow from both WK and Osak, although the temporal order of these admixture events remains unresolved (Fig. 2e, Fig. S5d-f).

Introgressed chromosomal regions from EJ to Osak were assessed using *F_d_* statistics (EJ, WK; Osak, HSOK), calculated within sliding windows of 50 consecutive single-nucleotide variants (SNVs) along the chromosomes. The *F_d_*values were shifted toward the positive direction across the genome (Fig. S6), consistent with the *D* statistic results (Fig. 2b). This result indicates that the *F_d_* analysis is robust, even when an inbred individual (HSOK) is used as the outgroup. We interpreted the top 0.5% of windows in the genome-wide *F_d_* distribution as candidate introgression regions. Outliers of *F_d_* were detected on chromosomes, 1, 3, 4, 5, 6, 7, 12, 17, 18, 19 and 21 (Fig. 2f, Table S7, S8). To confirm the possible gene functions of 12 annotated genes on the introgressed regions, we identified ortholog groups and performed the GO term enrichment analysis (Tables S9), based on reference genome annotations for *O. latipes* (HdrR strain, ASM223467v1), and *O. sakaizumii* (HNI strain, ASM223471v1). The GO analysis indicated an enrichment of genes involved in odor sensing mediated by trace amine-associated receptors (GO:0001594 trace-amine receptor activity: *FDR* = 4.1 ✕10^−12^, *Fold enrichment* = 280.95, *nGenes* = 7) (Tables S8–S9); however, these genes are physically clustered on chromosome 21. In addition, an introgressed region on chromosome 6 was overlapped with two egg envelope genes, *1-sf* and *choriogenin H* (GO:0035805 egg coat: *FDR* = 0.0050, *Fold enrichment* = 129.67, *nGenes* = 2), which are homologs of zona pellucida genes in mammals and essential to produce egg envelopes in *O. latipes* (Murata and Kinoshita 2022).

### Selection signatures in Oryzias sakaizumii

To explore genetic regions with signatures of hard selective sweep in *O. sakaizumii* (Fig. S1d), we calculated the integrated haplotype score (iHS) for SNPs with minor allele frequencies greater than 5% in 40 individuals of *O. sakaizumii* (Fig. 3, Table 1, Table S10). We extracted the top 0.25% and bottom 0.25% of the empirical *Z*-transformed iHS distribution (986 SNPs, Table S10). Of these, 61 SNPs were contained in annotated genes (Table 1). In those SNPs, phosphate-binding genes tended to be enriched (GO:0032266 phosphatidylinositol-3-phosphate binding: *FDR* = 0.0250, *Fold enrichment* = 54.80, *nGenes* = 3, Table S9). SNPs with outlier iHS values were located on some genes whose molecular functions have been well studied in fishes, such as *cd302* (Natural immunity, Brown et al. 2016, Zhang et al. 2022), *BDKRB2* (Cardiovascular system development and locomotor ability, Emanueli 1999), *hoxa13a* (Fin formation and soft fin ray length, Nakamura et al. 2016, Corcoran et al. 2026), and *ar* (androgen receptor beta, Ogino et al. 2023).

**Figure 3.**
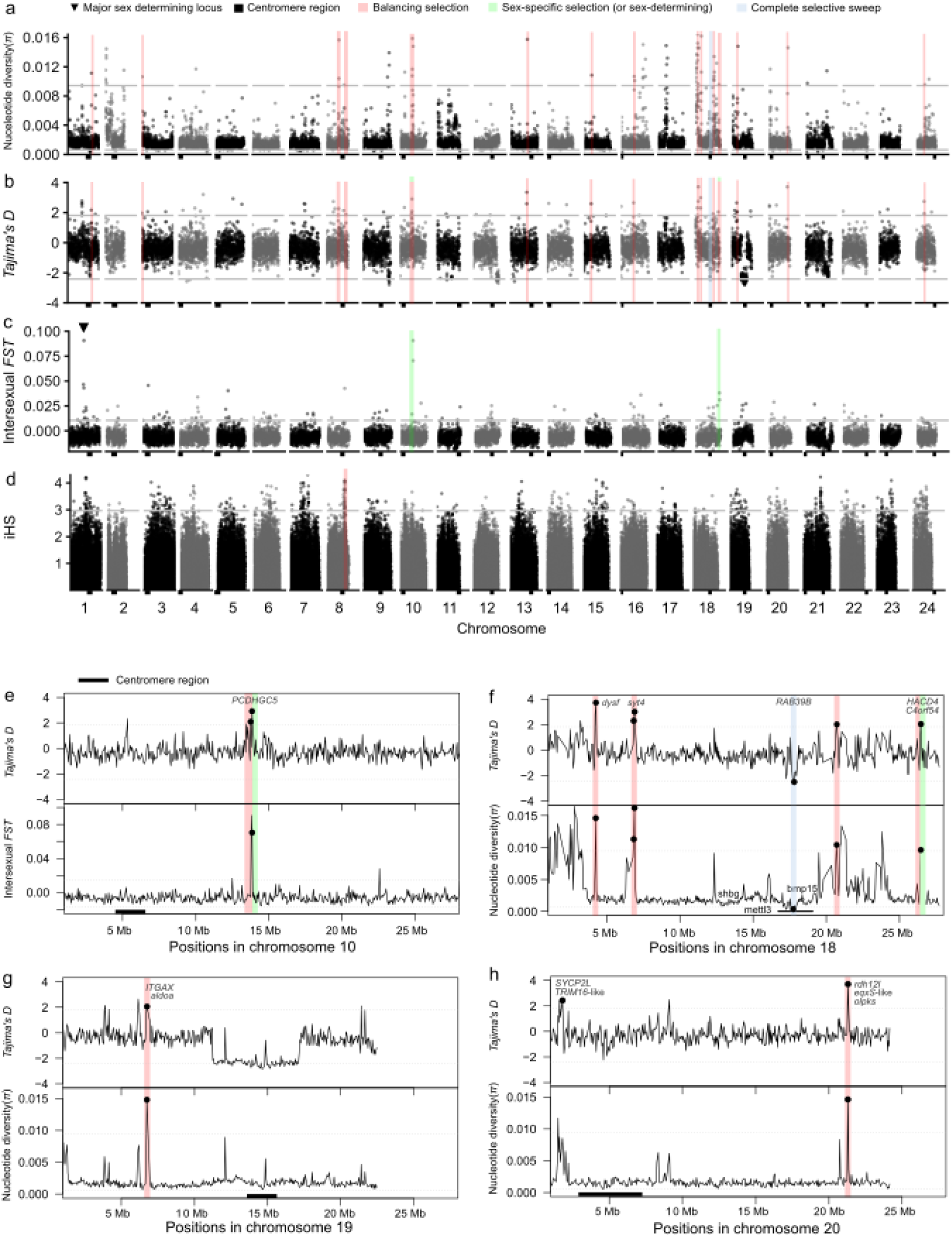
(a–c) Genome-wide selection scans using non-overlapping 50,000 bp windows across all 24 chromosomes of wild *O. sakaizumii*, showing nucleotide diversity π (a), *Tajima’s D* (b), and intersexual *FST* (c). Dotted lines indicate significance thresholds based on genome-wide distributions. (d) Integrated haplotype scores of 197253 SNPs across all 24 chromosomes. Each dot represents an absolute Z-transformed iHS value. Gray regions indicate genomic intervals with selection signatures in *O. sakaizumii*. (e-h) Selection signatures in chromosome 10, 18, 19, 20, respectively. Gene annotation information was retrieved from Ensembl 109, *Oryzias latipes* (HdrR strain, ASM223467v1). Black bars on X-axis in each panel represented the 1 Mb region near the inferred centromere region.

**Table 1.**
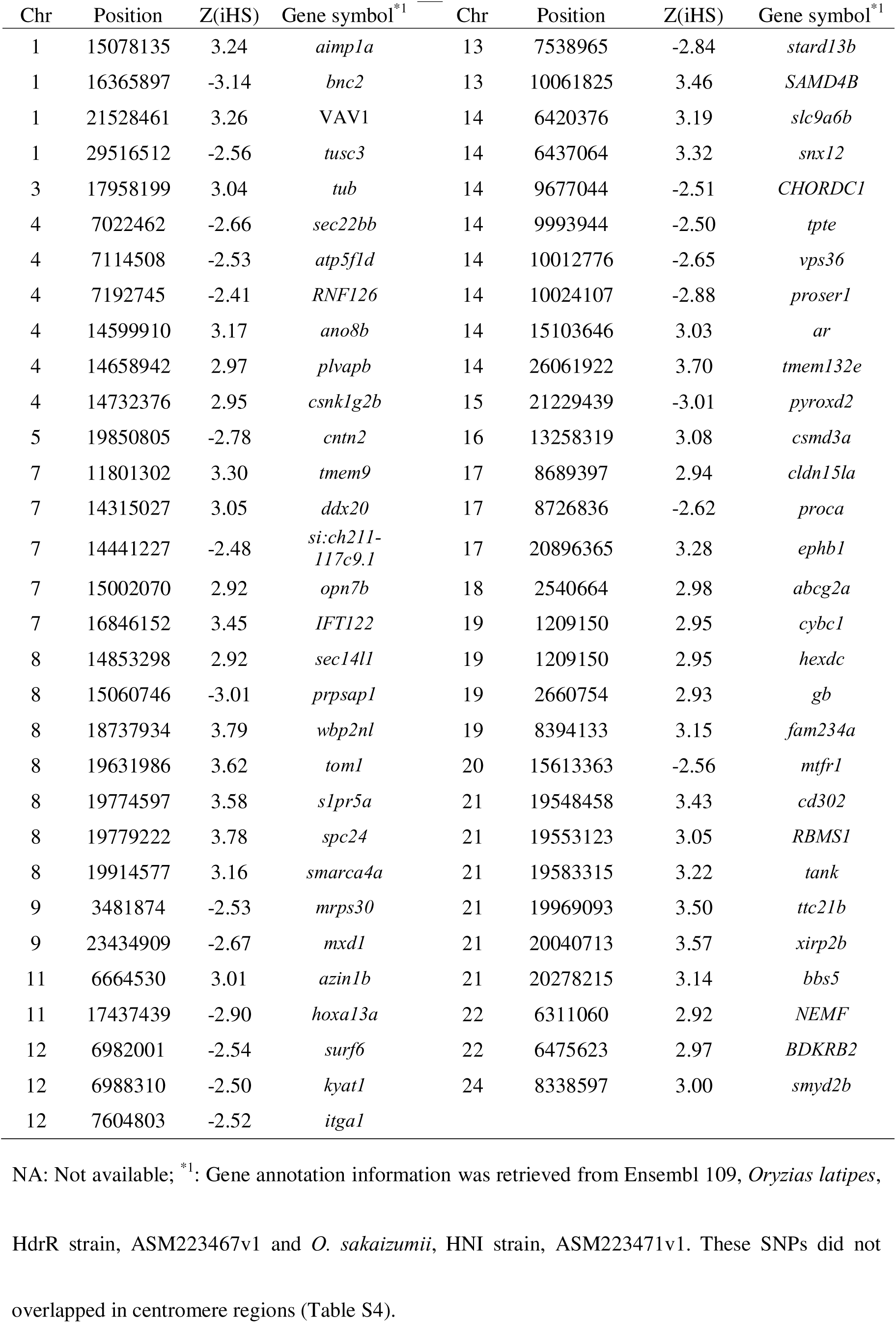
Genes containing SNPs with a selective sweep based on integrated haplotype score (iHS) in *O. sakaizumii*.

We also examined nucleotide diversity π, *Tajima’s D*, and genetic differentiation between sexes (intersexual *FST*) in non-overlapping 50-kb windows to identify signatures of selection (Fig. 3, Fig. S7, Table 2, Table S8). Based on outliers (top or bottom 0.5%) of the empirical distribution of the statistics (Fig. S1), we extracted (1) windows with high π and high *Tajima’s D* as candidates for balancing selection, (2) windows with low π and low *Tajima’s D* as candidates for complete selective sweep, and (3) windows with high intersexual *FST* as candidates for loci associated with sex-determination or sex-specific selection, where different alleles might be more advantageous for survival in each sex (Fig. S1). We observed overlaps between (1) and (3).

**Table 2.** Outlier regions of genetic diversity based on nucleotide diversity π, *Tajima’s D*, intersexual *FST*, and integrated haplotype score (iHS) within *O. sakaizumii*.

| High genetic diversity regions by genome scan |  |  |  |  |  |  | Integrated haplotype score (iHS) |  |  |
| --- | --- | --- | --- | --- | --- | --- | --- | --- | --- |
| Chr | Start | End (bp) | $\pi$ | *Tajima's D* | *Intersexual FST* | $F_d$ | Outlier SNPs | Max \|iHS\| | Max \|iHS\| position |
| 1 | 26000001 | 26050000 | **0.011\*** | **2.1709\*** | 0.0000 | 0.3465 | 0 | 1.73 | 26040924 |
| 3 | 1 | 50000 | **0.011\*** | **1.974\*** | -0.0010 | 0.0491 | NA | NA | NA |
| 8 | 14350001 | 14400000 | **0.016\*** | **2.267\*** | -0.005 | -0.111 | 0 | 1.39 | 14382920 |
| 8 | 19750001 | 19800000 | **0.010\*** | **2.139\*** | -0.0121 | 0.0324 | 34 | 4.14 | 19779802 |
| 10 | 13750001 | 13800000 | **0.012\*** | **2.086\*** | -0.006 | -0.028 | 0 | 2.01 | 13798063 |
| 10 | 13850001 | 13900000 | **0.016\*** | **2.908\*** | **0.070\*** | -0.019 | 0 | 2.31 | 13878057 |
| 13 | 18550001 | 18600000 | **0.016\*** | **3.363\*** | -0.013 | -0.152 | 0 | 1.80 | 18596925 |
| 15 | 8050001 | 8100000 | **0.011\*** | **2.583\*** | -0.009 | 0.029 | 0 | 1.90 | 8097458 |
| 16 | 14400001 | 14450000 | **0.011\*** | **2.649\*** | -0.012 | 0.017 | 0 | 1.66 | 14445917 |
| 18 | 4250001 | 4300000 | **0.015\*** | **3.719\*** | -0.011 | -0.083 | 0 | 1.70 | 4278760 |
| 18 | 6850001 | 6900000 | **0.011\*** | **2.302\*** | -0.011 | -0.090 | 0 | 2.23 | 6884357 |
| 18 | 6900001 | 6950000 | **0.016\*** | **2.993\*** | -0.002 | 0.004 | 0 | 2.43 | 6944225 |
| 18 | 17800001 | 17850000 | <sup>a</sup> **0.0003<sup>†</sup>** | **-2.486<sup>†</sup>** | 0.001 | -0.020 | NA | NA | NA |
| 18 | 20700001 | 20750000 | **0.0103\*** | **1.913\*** | -0.008 | -0.169 | 0 | 1.72 | 21303987 |
| 18 | 26450001 | 26500000 | **0.010\*** | **2.049\*** | **0.031\*** | -0.062 | 0 | 1.51 | 26489032 |
| 19 | 6750001 | 6800000 | **0.015\*** | **2.067\*** | -0.007 | 0.017 | 0 | 1.87 | 6781462 |
| 20 | 21300001 | 21350000 | **0.015\*** | **3.719\*** | -0.005 | 0.106 | 0 | 1.71 | 21303987 |
| 24 | 9500001 | 9550000 | **0.009\*** | **2.738\*** | -0.010 | -0.066 | 0 | 1.01 | 9511329 |

Bold texts with \* and † represented top 0.5% and bottom 0.5%, respectively, of the empirical distribution of the statistics; a: window overlapped in the
centromere regions. NA (not available): No SNPs to calculate iHS values in the window.

Genomic regions with high π and high *Tajima’s D* exhibited an enrichment of genes involved in “cell-cell adhesion” (GO:0098609, cell-cell adhesion molecules: *FDR* = 3.0✕10^−10^, *Fold enrichment* = 18.23, *nGenes* = 12; Tables S8, S9, significance level: *FDR* < 0.01) and “calcium ion binding” (GO:0005509, calcium ion binding: *FDR* = 0.0022, *Fold enrichment* = 7.811, *nGenes* = 7; Tables S8, S9). The genes associated with “cell-cell adhesion” included protocadherin alpha family and Integrin alpha-M-like (Table S8). The genes associated with “calcium ion binding” included synaptotagmin4 *syt4* (Tables S9, S11), which functions in synaptic development and dopamine transport (Delignat-Lavaud et al. 2022; Kim et al. 2024). Of the regions showing high π and high *Tajima’s D*, two regions on chromosomes 10 and 18 also exhibited high intersexual *FST* (Fig. 3, Table 2, Table S8).

In the search for signatures of selection, false positives are likely to occur in regions with low recombination rates, such as the near centromere regions (Stevison and McGaugh 2020, Peñalba and Wolf 2020). Based on the centromere region of *O. latipes*, Hd-rR (Ichikawa et al. 2017; Furuyama et al. 2019; Ansai et al. 2023, Table S4), we estimated the centromere region of *O. sakaizumii* and examined the overlap between the signatures of selection and the 1 Mb region near the centromere (Table 2, Table S10). Of the 984 SNPs with outlier iHS values, 58 SNPs were located within the 1 Mb region near the centromere regions (Table S10). None of the regions with balancing selection or sex-specific selection detected in our analyses were near the centromere (Table 2). The region with the signature of low genetic diversity (chr18:17.71-18.07M) overlapped with the centromere region (Fig. 3f, Table 2), indicating the signature may have resulted from low recombination and genetic drift, rather than complete selective sweep.

## Discussion

### Population structure shaped by isolation and introgression

The admixture analyses revealed a complex population history involving isolation and secondary contact within the *O. latipes* species complex (Fig. 2e, Fig. S5). Our results indicate that admixture occurred between *O. sakaizumii* and part of the EJ population of *O. latipes* (Fig. 2). Because the four collection sites of *O. sakaizumii* (i.e., Tsuruga, Niigata, Aomori, and Kitatsugaru), which are located near the southern and northern edges of the species’ distribution, exhibited consistent admixture signals in *D* statistics (Fig. 2b–d, Table S6), this admixture event likely occurred in the ancestral lineage of *O. sakaizumii*. Based on estimated divergence times, the ancestral lineage of *O. sakaizumii* dates between 3.06 × 10^4^ and 4.80 × 10^5^ generations ago (Fig. S4). Notably, *O. sakaizumii* is estimated to have dispersed over a range of 700 km within eastern mainland Japan during the past 30,000 years (Fig. 1, Fig. S4), suggesting that wild medaka populations possess substantial dispersal ability during initial colonization (Figs. 1, 2, Fig. S4, see also, Katsumura et al. 2019). Long-distance dispersal and/or admixture after secondary contact has also been reported in congeneric species distributed in the Indonesian islands (Sutra et al. 2019; Horoiwa et al. 2021; Montenegro et al. 2022; Flury et al. 2023). The admixture graph also revealed complex secondary contacts among *O. latipes* populations (Fig. 2e, Fig. S5), supporting the existence of gene flow among geographic populations.

If the ranges of geographic populations in Japanese archipelago are not constrained by dispersal ability, it remains unclear which factors have maintained the distinct geographic population structure observed today (Fig. 1–2). Based on the genetic variation in juvenile growth rates and female reproductive traits, latitudinal adaptation to climatic environments has been proposed in *O. sakaizumii* and *O. latipes* (Kawajiri et al. 2009; Shinomiya et al. 2023; Fujimoto et al. 2024, 2026). Thus, natural selection and life-history adaptation to local climate may limit gene flow between geographic populations. Similar patterns of genetic differentiation associated with local adaptation have been reported for other widely distributed fish species distributed across the Japanese archipelago using genome sequence data (e.g., Hirase et al. 2025).

### Possible target gene of selection

Previous selection scans in medaka fish in Japan have focused only on a single collection site of *O. latipes* (Spivakov et al. 2014; Fitzgerald et al. 2022). Based on the geographic location of collection site, the population in previous studies is considered to correspond to the Eastern Japan (EJ) population of *O. latipes* (Fig. 1, Katsumura et al. 2019). In this study, we investigated chromosomal regions exhibiting signatures of natural or sexual selection in wild *O. sakaizumii* for the first time. In the scan for selective sweeps by iHS, 61 SNPs located within genes were detected (Table 1).

Some iHS outliers are located on genes associated with the male-specific morphology and courtship behaviors (Table 1, Table S11). For example, androgen receptor beta is known to regulate the expression of male-specific traits in *O. latipes* (Ogino et al. 2023). In the zebrafish, *hoxa13a* affects fin formation and soft fin ray length in concert with its paralogs, *hoxa13b* and *hoxd13a* (Corcoran et al. 2026). In addition, iHS outliers on chromosome 7 are sporadically distributed along 11.73-15.60Mb (Fig. 3, Table S10). This region falls within a locus associated with fin morphology (Chr7:9.86-17.10Mb) in a genome-wide association study of ornamental medaka strains (Kon et al. 2026). Medaka exhibits sexual dimorphisms in anal and dorsal fin lengths, and males with longer fin lengths are more readily accepted by females in *O. sakaizumii* (Fujimoto et al. 2014, 2021). Sexual selection via female mate choice may lead to selective sweeps at loci associated with fin morphology through increased male reproductive success in wild populations.

We also identified distinct signals of introgression from the EJ population into *O. sakaizumii* on several chromosomes (Fig. 2, Table S7, S11). In the chr6:18.15-18.18Mb region, we confirmed that eight individuals of *O. sakaisumii* are nested within *O. latipes*, while EJ and *O. sakaizumii* together form a monophyletic clade (Fig. S8). The topology supports the introgressed allele from *O. latipes* are nearly fixed in wild *O. sakaisumii* populations. Genetic variation in *1-sf*, *choriogenin h*, and *cpt1b*, may have fitness effects through egg formation or lipid metabolism during female reproduction. However, it remains unclear whether phenotypic differences in egg characteristics among the introgressed allele of EJ (Osak) and the alleles found in WJ and WK populations. Comparative analyses of phenotypes across *O. latipes* subpopulations will be essential for evaluating the functional significance of the introgressed region.

Our selection scan for high π and high *Tajima’s D* regions identified candidate genetic regions under balancing selection in wild *O. sakaizumii* (Fig. 3, Table 2, Table S11). Within these genomic regions, the *synaptotagmin 4* (*syt4*) gene emerges as a candidate influencing mating behavior. *syt4* functions in synaptic development and dopamine transport (Delignat-Lavaud et al. 2022; Kim et al. 2024). In humans and mice, *syt4* gene expression affects individual personality traits, such as prosocial activity and depressive symptoms (Delignat-Lavaud et al. 2022; Kim et al. 2024). In avian species, such as *Ficedula albicollis* and *F. hypoleuca*, brain expression levels of *syt4* differ between species. Since both species can learn songs into adulthood, divergent gene expression levels of *syt4* in adults may contribute to species-specific male song repertoires (Wheatcroft et al. 2025).

Notably, the locus including *syt4* (chr18:6.85–6.95M) located in a QTL associated with a female mating behavior, spawning latency (chr18:3.71-12.64M, Fujimoto et al. 2024, Table S11). Spawning latency in the medaka females is an index of mate choosiness, i.e., how amount a female rejects the male courtship and copulation until egg spawning (Okuyama et al. 2014; Fujimoto et al. 2015). In addition, the high genetic diversity in the region could not be attributed to genetic differentiation between the sexes (Table 2), sex-determining mechanism (Kitano et al. 2023) or sex-specific selection is an unlikely to be the cause of observed high genetic diversity in the locus. Based on these features of the population genetic statistics in the locus, we hypothesize that sexual selection related to female mating behavior is the cause of balancing selection.

Previously reported loci of QTLs in medaka are located on regions spanning several megabases on chromosomes, with each locus typically containing 100–200 genes (e.g., Kawajiri et al. 2014, Fujimoto et al. 2024). Therefore, further detailed genetic analysis is needed to narrow down the locus to the causal polymorphic sites. Recent efforts to identify genes associated with mate choice behavior have employed not only QTL mapping but also candidate gene approaches, including genome editing to verify gene-phenotype relationships (Rossi et al. 2020, 2024; Ansai et al. 2021). Although our selection scans do not provide direct evidence of the association between genotypes and phenotypes, it can be contributed to identifying candidates for the causal polymorphic sites.

### Sexual selection in female mating behavior

Female medaka fish spawn small batches of eggs repeatedly throughout the reproductive season via external fertilization. This reproductive strategy may involve an evolutionary trade-off between female mate assessment and fitness. While larger, dominant males are preferred by females and mate frequently (Howard et al. 1998; Fujimoto et al. 2015), such high mating frequency in a male lead to sperm depletion which, reduces fertilization success (Weir and Grant 2010; Kondo et al. 2025a). Since daily spawning activity peaks in the morning (Kondo et al. 2025b), females with longer spawning latencies are likely to become mate later part of daily spawning. Because sperm-depleted males continue to court females (Kondo et al. 2025a), if females cannot precisely discriminate against depleted males, prolonged assessment increases the risk of mating with depleted males. Female spawning latency evolves under a trade-off between the potential benefits of mate choice and the costs of reduced fertilization by sperm depletion (e.g., Pomiankowski and Iwasa 1998).

Genetic polymorphism maintained by sexual selection provides insight into how variation persists within populations and drives diversification of sexual traits between geographic populations or species. Balancing selection typically prevents any single optimal allele from reaching fixation (Hedrick 2007; Fijarczyk and Babik 2015), thereby maintaining a reservoir of standing genetic variation. This variation may serve as raw material for adaptive evolution in different environments. For example, *O. latipes* WK females exhibit a significantly longer average spawning latency than *O. sakaizumii*, regardless of the male’s origin (Fujimoto et al. 2015). The differences in population average of spawning latency suggest that mating strategies have evolved in response to the local ecological conditions, such as sex ratio (Kappler et al. 2023; Fujimoto et al. 2024) and the availability of potential mates (Kokko and Johnstone 2002).

### Concluding remarks

We performed whole-genome resequencing of wild medaka populations in Japan to reconstruct past admixture scenarios in the eastern part of mainland Japan. Medaka and its relatives provide an excellent model system for investigating how genetic polymorphisms shape phenotypes and their evolutionary consequences under complex admixture histories. We detected candidates of introgressed loci in wild populations of *O. sakaizumii* from *O. latipes*. In addition, we examined genomic regions of high genetic diversity and high *Tajima’s D* in wild *O. sakaizumii* to detect signatures of balancing selection. Since a candidate region under balancing selection overlaps with a QTL associated with female mating behavior, our findings suggest that sexual selection contributes to the maintenance of genetic polymorphisms.

## Supporting information

Suppelementary Figures

## Acknowledgments

The authors thank Mizuki Horoiwa and Kazunori Yamahira for discussions and feedback, and Jun Kitano for advice on data analysis. Takahiro Irie and Ryo Kakioka for assistance with wild fish collection, and Yukuto Sato for help with molecular experiments and computing resources. We thank the anonymous reviewers for their helpful feedback. Molecular experiments and computational analyses were conducted using instruments at the Center for Research Advancement and Collaboration and the Research Laboratory Center, Faculty of Medicine, University of the Ryukyus. The authors acknowledge the use of Gemini (https://gemini.google.com/) and DeepL (https://www.deepl.com/) to English language editing. The authors thank FORTE Science Communications (https://www.forte-science.co.jp/) for English language editing. This study was supported in part by the Spatiotemporal Genomics Project and advanced medical research grant in the University of the Ryukyus. This study was also supported by the JSPS KAKENHI Grant to SF (JSPS KAKENHI Grant Numbers JP19K16232 and 24K09610), as well as a Nakatsuji Foresight Foundation Research Grant to SF.

## Ethics statement

All methods related to the handling of live fish were conducted in accordance with the Regulations for Animal Experiments at University of the Ryukyus. All experiments were approved by the Animal Care Ethics Committee of University of the Ryukyus (Approval numbers: R2019035, R2021012, and R2024022). All experimental methods are reported in accordance with ARRIVE guidelines.

## Data accessibility statement

Genetic data: Raw sequence reads are deposited in the DDBJ Sequence Read Archive (DRA) database (BioProject: PRJDB20255). Individual genotype data (VCF) are available on Dryad (https://datadryad.org/share/10.5061/dryad.59zw3r2mt, Not currently available). Linux scripts for population genetic analysis and custom R scripts are available on Github (https://github.com/shinFujimoto/popGenomeMedaka).

## Author contributions

SF and TM conceived and designed the study. SF, HK, IM, BKAS, MY, and TK collected specimens from wild populations. SF, TM, and RK contributed to securing funding for this research. SF, IM, and BKAS conducted the mating experiments. SF, HA, and TM performed molecular laboratory work. HA provided access to the instruments necessary for experiments and data analysis. SF, MM, and RK conducted population genetic analyses. SF and RK wrote the draft. All authors revised the manuscript and approved the final version for submission.

## References

Abdul-Rahman F, Tranchina D, Gresham D. 2021. Fluctuating environments maintain genetic diversity through neutral fitness effects and balancing selection. Mol Biol Evol. 38:4362–4375. 10.1093/molbev/msab173

Alexander DH, Novembre J, Lange K. 2009. Fast model-based estimation of ancestry in unrelated individuals. Genome Res, 19:1655–1664. 10.1101/gr.094052.109

Andersson M, Simmons LW. 2006. Sexual selection and mate choice. Trends Ecol Evol. 21:296–302. 10.1016/j.tree.2006.03.015

Ansai S, Mochida K, Fujimoto S, Mokodongan DF, Sumarto BKA, Masengi KWA, Hadiaty RK, Nagano AJ, Toyoda A, Naruse K, et al. 2021. Genome editing reveals fitness effects of a gene for sexual dichromatism in Sulawesian fishes. Nat Comm. 12:1–13. 10.1038/s41467-021-21697-0

Ansai S, Montenegro J, Masengi KW, Nagano AJ, Yamahira K, Kitano J. 2022. Diversity of sex chromosomes in Sulawesian medaka fishes. J Evol Biol. 35:1751–1764. 10.1111/jeb.14076

Audzijonyte A, Richards SA. 2018. The energetic cost of reproduction and its effect on optimal life-history strategies. Am Nat. 192:E150–E162. 10.1086/698655

Auwera V der, O’Connor BD. 2020. Genomics in the cloud: using Docker, GATK, and WDL in Terra. Sebastopol: O’Reilly Media.

Bissegger M, Laurentino TG, Roesti M, Berner D. 2020. Widespread intersex differentiation across the stickleback genome–The signature of sexually antagonistic selection? Mol Ecol. 29:262–271. 10.1111/mec.15255

Bonduriansky R, Chenoweth,SF. 2009. Intralocus sexual conflict. Trends Ecol Evol. 24:280–288. 10.1016/j.tree.2008.12.005

Cook DE, Andersen EC. 2017. VCF-kit: Assorted utilities for the variant call format. Bioinformatics. 33:1581–1582. 10.1093/bioinformatics/btx011

Cole JM, Scott CB, Johnson MM, Golightly PR, Carlson J, Ming MJ, Harpak A, Kirkpatrick M. 2024. The battle of the sexes in humans is highly polygenic. Proc. Natl. Acad. Sci. U.S.A. 121:e2412315121.

Coombe L, Kazemi P, Wong J, Birol I, Warren RL. 2024. Multi-genome synteny detection using minimizer graph mappings. bioRxiv, 2024-02, last accessed December 13, 2025. 10.1101/2024.02.07.579356

Corcoran J, Quigley H, Qu Q, Akimenko MA. 2026. Combined mutations of *hoxa13a*, *hoxa13b*, and *hoxd13a* lead to structural shifts in zebrafish soft fin rays providing insight into spiny ray evolution. PLoS genet, 22: e1012055. 10.1371/journal.pgen.1012055

Daimon M, Katsumura T, Sakamoto H, Ansai S, Takeuchi H. 2022. Mating experiences with the same partner enhanced mating activities of naïve male medaka fish. Scientific reports, 12(1), 19665. 10.1038/s41598-022-23871-w

Danecek P, Bonfield JK, Liddle J, Marshall J, Ohan V, Pollard MO, Whitwham A, Keane T, McCarthy SA, Davies RM. 2021. Twelve years of SAMtools and BCFtools. GigaScience. 10:1–4. 10.1093/gigascience/giab008

Delaneau O, Zagury J, Robinson MR, Marchini J, Dermitzakis E. 2019. Accurate, scalable and integrative haplotype estimation. Nat Comm. 10:5436. 10.1038/s41467-019-13225-y

Delignat-Lavaud BT, Ducrot C, Kouwenhoven W, Feller N, Trudeau LR. 2022. Implication of synaptotagmins 4 and 7 in activity-dependent somatodendritic dopamine release in the ventral midbrain. Open Biol. 12:210339. 10.1098/rsob.210339

Dong Z, Wang J, Chen G, Guo Y, Zhao N, Wang Z, Zhang B. 2024. A high-quality chromosome-level genome assembly of the Chinese medaka *Oryzias sinensis*. Sci Data. 11:6–11. 10.1038/s41597-024-03173-8

Emanueli C, Maestri R, Corradi D, Marchione R, Minasi A, Tozzi MG. 1999. Dilated and failing cardiomyopathy in bradykinin B2 receptor knockout mice. Circulation, 100:2359–2365. 10.1161/01.CIR.100.23.2359

Emms DM, Kelly S. 2019. OrthoFinder: Phylogenetic orthology inference for comparative genomics. Genome Biol. 20:1–14. 10.1186/s13059-019-1832-y

Excoffier L, Dupanloup I, Huerta-Sánchez E, Sousa VC, Foll M. 2013. Robust Demographic Inference from Genomic and SNP Data. PLoS Genet. 9:e1003905. 10.1371/journal.pgen.1003905

Ferrer-Admetlla A, Liang M, Korneliussen T, Nielsen R. 2014. On detecting incomplete soft or hard selective sweeps using haplotype structure. Mol Biol Evol. 31:1275–1291. 10.1093/molbev/msu077

Fijarczyk A, Babik W. 2015. Detecting balancing selection in genomes: Limits and prospects. Mol Ecol. 24:3529–3545. 10.1111/mec.13226

Fitzgerald T, Brettell I, Leger A, Wolf N, Kusminski N, Monahan J, Barton C, Herder C, Aadepu N, Gierten, et al. 2022. The medaka inbred Kiyosu-Karlsruhe (MIKK) panel. Genome Biol. 23:59. 10.1186/s13059-022-02623-z

Flury JM, Meusemann K, Martin S, Hilgers L, Spanke T, Böhne A, Herder F, Mokodongan DF, Altmüller J, Wowor D, et al. 2023. Potential contribution of ancient introgression to the evolution of a derived reproductive strategy in ricefishes. Genome Biol Evol. 15:1–12. 10.1093/gbe/evad138

Fujimoto S, Kawajiri M, Kitano J, Yamahira K. 2014. Female mate preference for longer fins in medaka. Zool Sci, 31:703–708. 10.2108/zs140102

Fujimoto S, Miyake T, Yamahira K. 2015. Latitudinal variation in male competitiveness and female choosiness in a fish: are sexual selection pressures stronger at lower latitudes? Evol Biol. 42:75–87. 10.1007/s11692-014-9300-9

Fujimoto, S., Murase, I., Kobayashi, H., Sumarto, B. K., Yagi, M., & Yamahira, K. 2026. Shorter recruitment periods and higher fecundity in high-latitude populations of the *Oryzias latipes* species complex. Popl Ecol, 68, e70012. 10.1002/1438-390x.70012

Fujimoto S, Sumarto BKA, Murase I, Mokodongan DF, Myosho T, Yagi M, Ansai S, Kitano J, Takeda S, Yamahira K. 2024. Evolution of size-fecundity relationship in medaka fish from different latitudes. Mol Ecol. 33:e17578. 10.1111/mec.17578

Fujimoto S, Takeda S, Yagi M, Yamahira, K. 2021. Seasonal change in male reproductive investment of a fish. Env Biol Fish. 104:107–118. 10.1007/s10641-021-01059-x

Fujimoto S, Yaguchi H, Myosho T, Aoyama H, Sato Y, Kimura R. 2022. Population admixtures in medaka inferred by multiple arbitrary amplicon sequencing. Sci Rep. 12:19989. 10.1038/s41598-022-24498-7

Ge SX, Jung D, Jung D, Yao R. 2020. ShinyGO: A graphical gene-set enrichment tool for animals and plants. Bioinformatics. 36:2628–2629. 10.1093/bioinformatics/btz931

Gel B, Díez-Villanueva A, Serra E, Buschbeck M, Peinado MA, Malinverni R. 2016. regioneR: an R/Bioconductor package for the association analysis of genomic regions based on permutation tests. Bioinformatics. 32:289–291. 10.1093/bioinformatics/btv562

Howard RD, Martens RS, Innis SA, Drnevich JM, Hale J. 1998. Mate choice and mate competition influence male body size in Japanese medaka. Anim Behav, 55:1151–1163. 10.1006/anbe.1997.0682

Hedrick PW. 1999. Antagonistic pleiotropy and genetic polymorphism: a perspective. Heredity. 82:126–133. 10.1046/j.1365-2540.1999.00440.x

Hedrick PW. 2007. Balancing selection. Curr Biol. 17:230–231.

Hinrichs AS, Karolchik D, Baertsch R, Barber GP, Bejerano G, Clawson H, Diekhans M, Furey TS, Harte RA, Hsu F, et al. 2006. The UCSC Genome Browser Database: update 2006. Nucleic Acids Res. 34:590–598. 10.1093/nar/gkj144

Hirase S, Nagano AJ, Nohara K, Kikuchi K, Kokita T. 2025. Phenotypic and genomic signatures of latitudinal local adaptation along with prevailing ocean current in a coastal goby. Mol Ecol. 34: e17599. 10.1111/mec.17599

Horoiwa M, Mandagi IF, Sutra N, Montenegro J, Tantu FY, Masengi KWA, Nagano AJ, Kusumi J, Yasuda N, Yamahira K. 2021. Mitochondrial introgression by ancient admixture between two distant lacustrine fishes in Sulawesi Island. PLoS ONE. 16:1–14. 10.1371/journal.pone.0245316

Huson DH, Bryant D. 2006. Application of phylogenetic networks in evolutionary studies. Mol Biol Evol. 23:254–267. 10.1093/molbev/msj030

Ichikawa K, Tomioka S, Suzuki Y, Nakamura R, Doi K, Yoshimura J, Kumagai M, Inoue Y, Uchida Y, Irie N, et al. 2017. Centromere evolution and CpG methylation during vertebrate speciation. Nat Comm. 8:1833. 10.1038/s41467-017-01982-7

Iwamatsu, T. 2006. The integrated book for the biology of the medaka. (In Japanese). Okayama, DAIGAKU KYOIKU SHUPPAN.

Furuyama M, Nagaoka H, Sato T, Sakaizumi M. 2019. Centromere localization in medaka fish based on half-tetrad analysis. GGS. 94:159–165.

Kappeler PM, Benhaiem S, Fichtel C, Fromhage L, Höner OP, Jennions MD et al. 2023. Sex roles and sex ratios in animals. Biol Rev, 98:462–480. 10.1111/brv.12915

Katsumura T, Oda S, Mitani H, Oota H. 2019. Medaka population genome structure and demographic history described via genotyping-by-sequencing. G3. 9:217–228. 10.1534/g3.118.200779

Katsumura T, Oda S, Nakagome S, Hanihara T, Kataoka H, Mitani H, Kawamura S, Oota H. 2014. Natural allelic variations of xenobiotic enzymes pleiotropically affect sexual dimorphism in *Oryzias latipes*. Proc R Soc B. 281:20142259. 10.1101/000661

Kawajiri M, Kokita T, Yamahira K. 2009. Heterochronic differences in fin development between latitudinal populations of the medaka *Oryzias latipes* (Actinopterygii: Adrianichthyidae). Biol J Linn Soc. 97:571–580.

Kawajiri M, Yoshida K, Fujimoto S, Mokodongan DF, Ravinet M, Kirkpatrick M, Yamahira K, Kitano J. 2014. Ontogenetic stage-specific quantitative trait loci contribute to divergence in developmental trajectories of sexually dimorphic fins between medaka populations. Mol Ecol. 23:5258–5275. 10.1111/mec.12933

Kent WJ. 2002. BLAT—the BLAST-like alignment tool. Genome res. 12:656–664. 10.1101/gr.229202

Kim J, Seol S, Kim TE, Lee J, Koo JW, Kang HJ. 2024. Synaptotagmin-4 induces anhedonic responses to chronic stress via BDNF signaling in the medial prefrontal cortex. Exp Mol Med, 56:329–343. 10.1038/s12276-024-01156-8

Kimura R, Fujimoto A, Tokunaga K, Ohashi J. 2007. A practical genome scan for population-specific strong selective sweeps that have reached fixation. PLoS one. 2:e286. 10.1371/journal.pone.0000286

Kitano J, Ansai S, Fujimoto S, Kakioka R, Sato M, Mandagi IF, Sumarto BKA, Yamahira K. 2023. A cryptic sex-linked locus revealed by the elimination of a master sex-determining locus in medaka fish. Am Nat, 202:231–240. 10.1086/724840

Kokko, H., & Johnstone, R. A. 2002. Why is mutual mate choice not the norm? Operational sex ratios, sex roles and the evolution of sexually dimorphic and monomorphic signalling. Philosophical Transactions of the Royal Society of London. Series B: Biological Sciences, 357(1419), 319–330.

Kon T, Tang R, Kon-Nanjo K, Tomihara S, Fushiki S, Fujii W et al. 2026. Genomic consequences of domestication and the diversification of body coloration and morphology in ornamental medaka strains. Mol Biol Evol, 43:msag021. 10.1093/molbev/msag021

Kondo Y, Kohda M, Awata S. 2025a. Male medaka continue to mate with females despite sperm depletion. R Soc Open Sci. 12:241668. 10.1098/rsos.241668

Kondo Y, Kobayashi R, Kobayashi Y, Awata S. 2025b. Temporal dynamics of courtship and spawning in medaka under laboratory conditions revealed by 24 h video monitoring. Sci Rep, 15:26576. 10.1038/s41598-025-11082-y

Korunes KL, Samuk K. 2021. pixy: Unbiased estimation of nucleotide diversity and divergence in the presence of missing data. Mol Ecol Res. 21:1359–1368. 10.1111/1755-0998.13326

Li H. 2013. Aligning sequence reads, clone sequences and assembly contigs with BWA-MEM. http://arxiv.org/abs/1303.3997, last accessed December 7, 2025

Maier R, Flegontov P, Flegontova O, Işıldak U, Changmai P, Reich D. 2023. On the limits of fitting complex models of population history to *f*-statistics. ELife. 12:1–62. 10.7554/eLife.85492

Malinsky M, Matschiner M, Svardal H. 2021. Dsuite - Fast D-statistics and related admixture evidence from VCF files. Mol Ecol Res. 21:584–595. 10.1111/1755-0998.13265

Malinsky M, Svardal H, Tyers AM, Miska EA, Genner MJ, Turner GF, Durbin R. 2018. Whole-genome sequences of Malawi cichlids reveal multiple radiations interconnected by gene flow. Nat Ecol Evol. 2:1940–1955. 10.1038/s41559-018-0717-x

Mank JE. 2017. Population genetics of sexual conflict in the genomic era. Nat Rev Genet. 18:721–730. 10.1038/nrg.2017.83

Martin SH, Davey JW, Jiggins CD. 2015. Evaluating the use of *ABBA-BABA* statistics to locate introgressed loci. Mol Biol Evol. 32:244–257. 10.1093/molbev/msu269

Mcbride RS, Somarakis S, Fitzhugh, GR, Albert A, Yaragina NA, Wuenschel MJ, Alonso-Fernández A, Basilone G. 2015. Energy acquisition and allocation to egg production in relation to fish reproductive strategies. Fish Fish. 16:23–57. 10.1111/faf.12043

Mérot C, Llaurens V, Normandeau E, Bernatchez L, Wellenreuther M. 2020. Balancing selection via life-history trade-offs maintains an inversion polymorphism in a seaweed fly. Nat Comm. 11:670. 10.1038/s41467-020-14479-7

Montenegro J, Fujimoto S, Ansai S, Nagano AJ, Sato M, Maeda Y, Tanaka R, Masengi KWA, Kimura R, Kitano J, et al. 2022. Genetic basis for the evolution of pelvic-fin brooding, a new mode of reproduction, in a Sulawesian fish. Mol Ecol. 31:3798–3811. 10.1111/mec.16555

Murata K, Kinoshita M. 2022. Targeted deletion of liver-expressed Choriogenin L results in the production of soft eggs and infertility in medaka, Oryzias latipes. Zool Lett, 8:1. 10.1186/s40851-021-00185-9

Nakamura T, Gehrke AR, Lemberg J, Szymaszek J, Shubin NH. 2016. Digits and fin rays share common developmental histories. Nature, 537(7619), 225–228. 10.1038/nature19322

Nishiike Y, Miyazoe D, Togawa R, Yokoyama K, Nakasone K, Miyata M, et al. 2021. Estrogen receptor 2b is the major determinant of sex-typical mating behavior and sexual preference in medaka. Curr Biol, 31, 1699–1710. 10.1016/j.cub.2021.01.089

Okuyama T, Yokoi S, Abe H, Isoe Y, Suehiro Y, Imada H, Tanaka M, Kawasaki T, Yuba S, Taniguchi Y, et al. 2014. A neural mechanism underlying mating preferences for familiar individuals in medaka fish. Science. 343:91–94. 10.1126/science.1244724

Ogino Y, Ansai S, Watanabe E, Yasugi M, Katayama Y, Sakamoto H, Okamoto K, Okubo K, Yamamoto Y, Hara I, et al. 2023. Evolutionary differentiation of androgen receptor is responsible for sexual characteristic development in a teleost fish. Nat comm. 14:1428.

Patterson N, Moorjani P, Luo Y, Mallick S, Rohland N, Zhan Y, Genschoreck T, Webster T, Reich D. 2012. Ancient admixture in human history. Genetics. 192:1065–1093. 10.1534/genetics.112.145037

Peñalba JV, Wolf JB. 2020. From molecules to populations: appreciating and estimating recombination rate variation. Nat Rev Genet. 21:476–492. 10.1038/s41576-020-0240-1

Peter BM. 2016. Admixture, population structure, and *f*-statistics. Genetics. 202:1485–1501. 10.1534/genetics.115.183913

Petr M, Vernot B, Kelso J. 2019. Admixr-R package for reproducible analyses using ADMIXTOOLS. Bioinformatics. 35:3194–3195. 10.1093/bioinformatics/btz030

Pomiankowski A, Iwasa Y. 1998. Runaway ornament diversity caused by Fisherian sexual selection. Proc Natl Acad Sci U.S.A. 95:5106–5111. 10.1073/pnas.95.9.5106

Pracana R, Priyam A, Levantis I, Nichols RA, Wurm Y. 2017. The fire ant social chromosome supergene variant Sb shows low diversity but high divergence from SB. Mol Ecol. 26:2864–2879. 10.1111/mec.14054

Radwan J, Engqvist L, Reinhold K. 2016. A paradox of genetic variance in epigamic traits: beyond “good genes” view of sexual selection. Evol Biol. 43:267–275. 10.1007/s11692-015-9359-y

Rossi M, Hausmann AE, Alcami P, Moest M, Roussou R, van Belleghem SM, Wright DS, Kuo CY, Lozano-Urrego D, Maulana A, et al. (2024). Adaptive introgression of a visual preference gene. Science. 383:1368–1373. 10.1126/science.adj9201

Rossi M, Hausmann AE, Thurman TJ, Montgomery SH, Papa R, Jiggins CD, McMillan WO, Merrill RM. 2020. Visual mate preference evolution during butterfly speciation is linked to neural processing genes. Nat Comm. 11:4763. 10.1038/s41467-020-18609-z

Sabeti PC, Reich DE, Higgins JM, Levine HZP, Richter DJ, Schaffner SF, Gabriel SB, Platko JV, Patterson NJ, McDonald GJ, et al. 2002. Detecting recent positive selection in the human genome from haplotype structure. Nature. 419:832–837.10.1038/nature01140

Schiffels S, Durbin R. 2014. Inferring human population size and separation history from multiple genome sequences. Nat Genet. 46:919–925. 10.1038/ng.3015

Schiffels S, Wang K. 2020. MSMC and MSMC2: the multiple sequentially Markovian coalescent In Statistical population genomics. In Methods in Molecular Biology. New York: Springer. p. 147–165.

Schliep KP. 2011. phangorn: Phylogenetic analysis in R. Bioinformatics. 27:592–593. 10.1093/bioinformatics/btq706

Shinomiya A, Adachi D, Shimmura T, Tanikawa M, Hiramatsu N, Ijiri S, Naruse K, Sakaizumi M, Yoshimura T. 2023. Variation in responses to photoperiods and temperatures in Japanese medaka from different latitudes. Zool Lett. 9:16. 10.1186/s40851-023-00215-8

Shinya M, Kimura T, Naruse K. 2023. High-speed system to generate congenic strains in medaka. GGS. 98:267–275. 10.1266/ggs.23-00075

Simmons LW, Lüpold S, Fitzpatrick JL. 2017. Evolutionary trade-off between secondary sexual traits and ejaculates. Trends Ecol Evol. 32:964–976. 10.1016/j.tree.2017.09.011

Soni V, Jensen JD. 2024. Temporal challenges in detecting balancing selection from population genomic data. G3. 14:jkae069. 10.1093/g3journal/jkae069

Spivakov M, Auer TO, Peravali R, Dunham I, Dolle D, Fujiyama A, Toyoda A, Aizu T, Minakuchi Y, Loosli F, et al. 2014. Genomic and phenotypic characterization of a wild medaka population: Towards the establishment of an isogenic population genetic resource in fish. G3. 4:433–445. 10.1534/g3.113.008722

Stevison, LS,McGaugh, SE. 2020. It’s time to stop sweeping recombination rate under the genome scan rug. Mol Ecol. 29: 4249–4253. 10.1111/mec.15690

Sutra N, Kusumi J, Montenegro J, Kobayashi H, Fujimoto S, Masengi KWA, Nagano AJ, Toyoda A, Matsunami M, Kimura R, Yamahira, K. 2019. Evidence for sympatric speciation in a Wallacean ancient lake. Evolution. 73:1898–1915. 10.1111/evo.13821

Szpiech ZA. 2024. selscan 2.0: scanning for sweeps in unphased data. Bioinformatics. 40:btae006. 10.1093/bioinformatics/btae006

Tajima F. 1989. Statistical method for testing the neutral mutation hypothesis by DNA polymorphism. Genetics. 123:585–595.

Takehana Y, Sakai M, Narita T, Sato T, Naruse K, Sakaizumi M. 2016. Origin of boundary populations in medaka (*Oryzias latipes* species complex). Zool Sci. 33:125–131. 10.2108/zs150144

Yassumoto TI, Nakatsukasa M, Nagano AJ, Yasugi M, Yoshimura T, Shinomiya A. 2020. Genetic analysis of body weight in wild populations of medaka fish from different latitudes. PloS One. 15:e0234803. 10.1371/journal.pone.0234803

Voight BF, Kudaravalli S, Wen X, Pritchard JK. 2006. A map of recent positive selection in the human genome. PLoS Biol. 4:e72. 10.1371/journal.pbio.0040072

Weir LK, Grant JWA. 2010. Courtship rate signals fertility in an externally fertilizing fish. Biol Lett, 6:727–731. 10.1098/rsbl.2010.0139

Wellenreuther M, Svensson EI, Hansson B. 2014. Sexual selection and genetic colour polymorphisms in animals. Mol Ecol. 23:5398–5414. 10.1111/mec.12935

Wheatcroft D, Backström N, Dutoit L, Mcfarlane SE, Mugal CF, Ålund M, Ellegren H, Qvarnström, A. 2025. Divergence in expression of a singing-related neuroplasticity gene in the brains of 2 Ficedula flycatchers and their hybrids. G3. 15:1–11. 10.1093/g3journal/jkae293

Wittmann MJ, Bergland AO, Feldman MW, Schmidt PS, Petrov DA. 2017. Seasonally fluctuating selection can maintain polymorphism at many loci via segregation lift. Proc Natl Acad Sci U.S.A. 114:E9932–E9941. 10.1073/pnas.1702994114

Zhang Z, Li Q, Huang Y, Jiang B, Li X, Huang M, et al. 2022. Molecular characterization of a novel C-type lectin receptors (*CD302*) in Nile tilapia (*Oreochromis niloticus*) and its functional analysis in host defense against bacterial infection. Aquaculture Reports, 27, 101405. 10.1016/j.aqrep.2022.101405

Zajitschek F, Connallon T. 2018. Antagonistic pleiotropy in species with separate sexes, and the maintenance of genetic variation in life-history traits and fitness. Evolution. 72:1306–1316. 10.1111/evo.13493

