## Supplementary material for "Genetic polymorphisms in a mate choice locus are maintained by balancing selection in a wild medaka population": Suppelementary Figures

Running title: Balancing selection in northern medaka

Shingo Fujimoto^†1^, Taijun Myosho ^2^, Hirozumi Kobayashi ^3^, Hiroaki Aoyama ^4^, Iki Murase ^5^, Bayu K. A. Sumarto ^6^, Mitsuharu Yagi ^7^, Taiga Kunishima ^8^, Masatoshi Matsunami ^9^, Ryosuke Kimura^†9^

^1^ Integrated Technology Center, University of the Ryukyus, Okinawa 903-2720, Japan.

^2^ Laboratory of Molecular Reproductive Biology, Institute for Environmental Sciences, University of Shizuoka, Shizuoka 422-8526, Japan

^3^ Natural History Museum and Institute Chiba, Chiba 260-8682, Japan

^4^ Research Facility Center, University of the Ryukyus, Okinawa 903-0213, Japan

^5^ School of Ocean and Earth Science, University of Southampton, Southampton, UK

^6^ Research Center for Conservation of Marine and Inland Water Resources, National Research and Innovation Agency, 16915, Indonesia

^7^ Graduate School of Fisheries and Environmental Sciences, Nagasaki University, Nagasaki 852-8521, Japan

^8^ Faculty of Agriculture, Setsunan University, 45-1 Nagaotoge, Hirakata, Osaka 573-0101, Japan

^9^ Graduate School of Medicine, University of the Ryukyus, Okinawa 903-2720, Japan

†: Corresponding authors

Ryosuke Kimura, Graduate School of Medicine, University of the Ryukyus, Okinawa 903-2720, Japan, Tel: +81-98-894-5166,

***Supporting information***

***Figure legends***

Figure. S1, Criteria and the numbers of extracted windows and SNPs in selection signatures

Figure S2. Results of ADMIXTURE analysis for *K* = 2–5. For the abbreviations of wild populations, see Tables S1 and S2.

Figure S3. (a) Geographic locations of collection sites. Solid circles and triangles on the map represent *Oryzias sakaizumii* and *Oryzias latipes*, respectively. Colors correspond to the clusters in the ADMIXTURE analysis (*K*= 4). (b) Neighbor-joining tree based on whole-genome sequences, excluding individuals with multiple ancestry in the ADMIXTURE analysis (*K* = 4). Excluded sites and individuals (Tng, Nss, Oit, Nck, and Nmg) were represented site names with underlined in the map.

Figure S4. (a, b) Relative cross-coalescence rates between four populations (a), and between collection sites in *O. sakaizumii* (b), calculated by MSMC2 based on four individuals (eight haplotypes). (c, d) Average divergence times in generations between populations (c) and schematic representation of population divergence (d).

Figure S5. (a–f) Admixture graphs that fit the data. Branch lengths (*F_2_* drift distances) are shown as solid lines, and admixture events are shown as dotted lines with mixture proportions indicated. (e) Introgressive regions identified by a genomic scan of *F_d_* statistics in non-overlapping 50-SNV windows.

Figure S6. Signature of introgressive regions represented with positive *F_d_* values in the combination of (a) EJ, WK, Osak, HSOK and (b) EJ, WJ, Osak, HSOK. *Oryzias sakaizumii*: Osak; *Oryzias latipes*: Western Kyushu (WK), Western Japan (WJ), and Eastern Japan (EJ).

Figure. S7, Nucleotide diversity, *Tajima’s D,* and Intersexual *FST* on chromosomes with signature of selection.

Figure. S8. Neighbor joining tree of introgressed locus on chromosome 6. *Oryzias sakaizumii*: Osak; *Oryzias latipes*: Western Kyushu (WK), Western Japan (WJ), and Eastern Japan (EJ)

***Table titles***

Table S1. Collection sites of samples of *Oryzias* *latipes* and *O. sakaizumii*.

Table S2. Sequence summary of individuals included in this study.

Table S3. One-to-one ortholog lists between the HdrR reference genome (ASM223467v1) and the HNI reference genome (ASM223471v1), based Ensembl release 109 annotations and Orthofinder results.

Table S4. Inferred position of centromere region in *O. sakaizumii*, HNI reference genome sequence (ASM223471v1). Physical positions were inferred based on the synteny block positions of HNI and HdrR reference sequence using ntSynt.

Table S5. Cross-validation (CV) errors of ADMIXTURE for *K* = 1 to 10.

Table S6. *Patterson’s D* statistics for five population combinations, *D* (X1, X2; Osak, HSOK), where X1 and X2 are WK, WJ, or EJ.

Table S7. Introgressed chromosomal regions in *O. sakaizumii* estimated by outliers *F_d_* (EJ, WK; Osak, HSOK) and signature of hard selective sweep shown by integrated haplotype score.

Table S8. Gene symbols in candidate regions of introgression and/or selection signatures. Annotation information was based on the Ensembl release 109, *Oryzias latipes* HdrR strain (ASM223467v1) and *O. sakaizumii* HNI strain (ASM223471v1).

Table S9. Top 20 gene ontology (GO) terms from enrichment analysis using ShinyGO ver. 0.82, sorted by ascending enrichment false discovery rate (FDR) (Significance threshold, *: *FDR* < 0.01; a: 0.01 < *FDR* < 0.05; N.S.: 0.05 < *FDR*).

Table S10. Results of integrated haplotype score in *Oryzias sakaizumii* calculated by selscan. ihh_0 and ihh_1 values corresponded to the reference allele and the alternative allele.

Table S11. Annotated genes located in regions with selection signatures and phenotypic associations identified in previous studies.

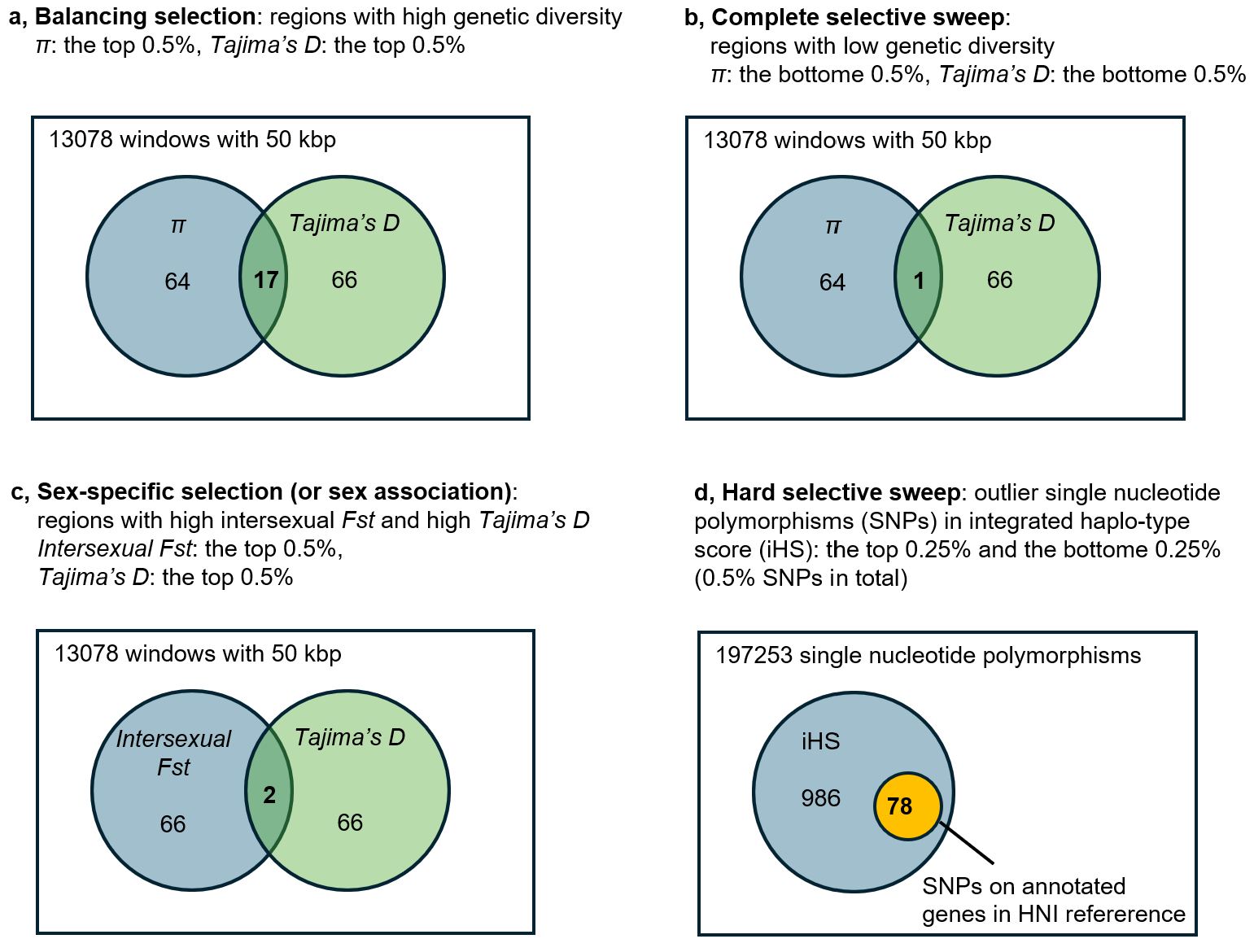

Figure. S1, Criteria and the numbers of extracted windows and SNPs in selection signatures

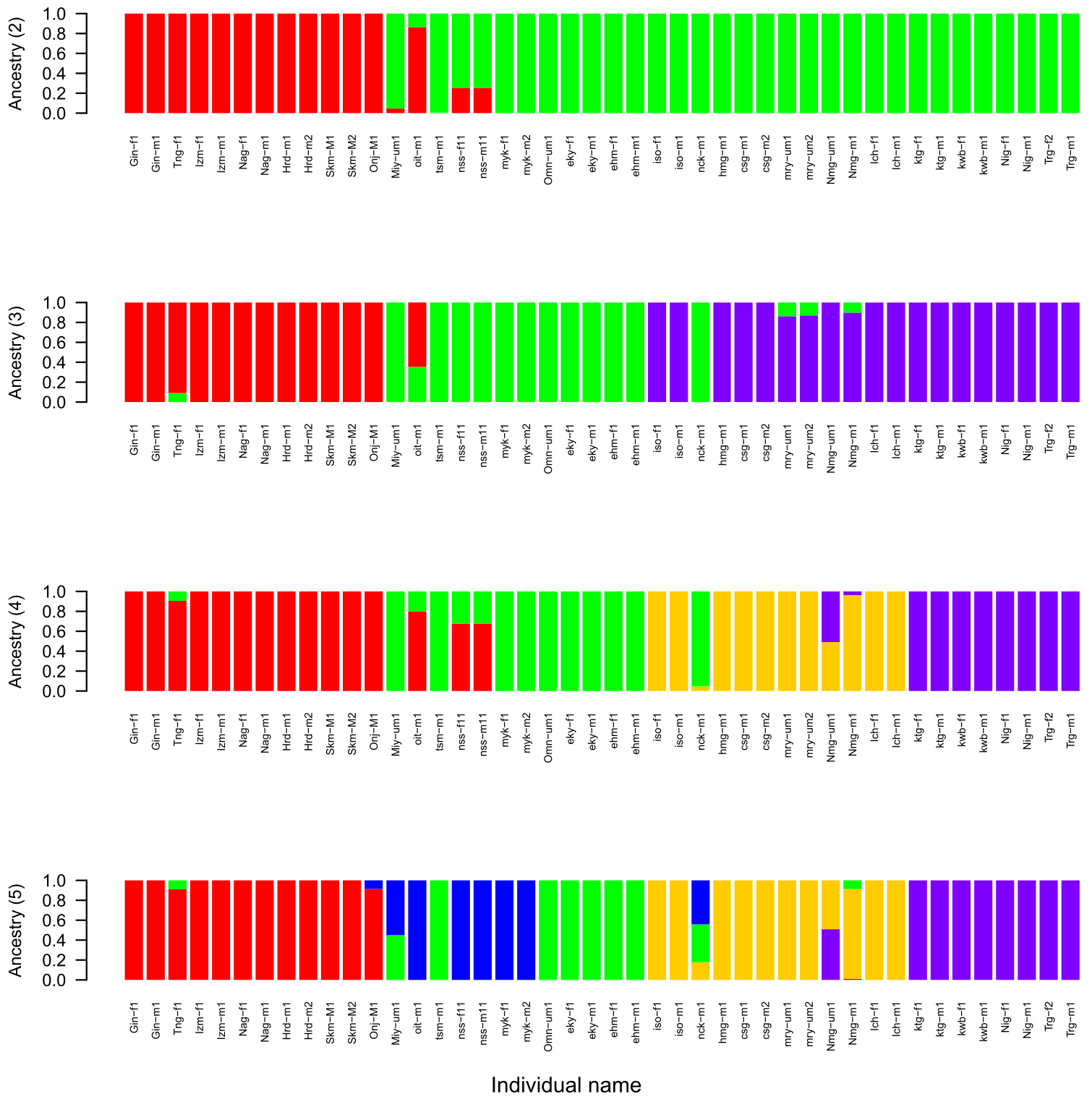

*O. sakaizumii*

Figure S2. Results of ADMIXTURE analysis for *K* = 2–5. For the abbreviations of wild populations, see Tables S1 and S2.

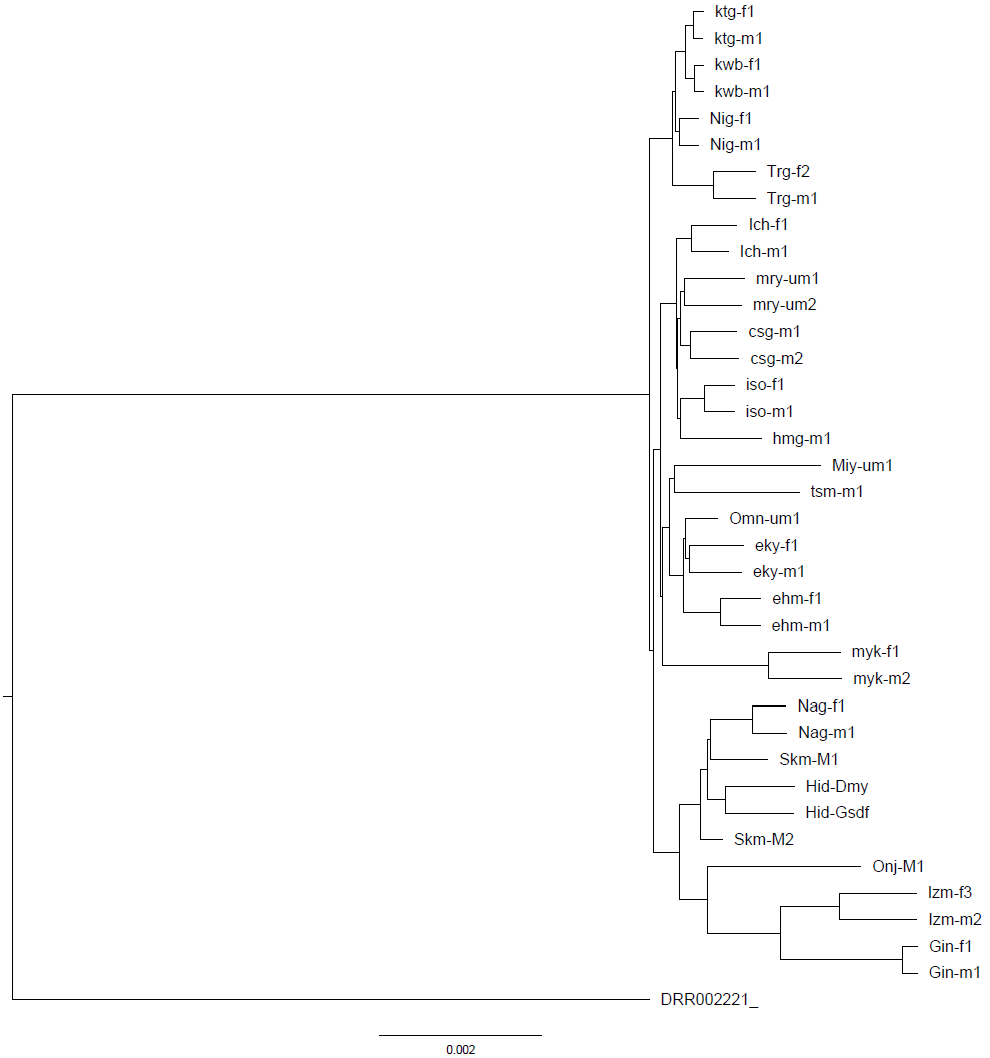

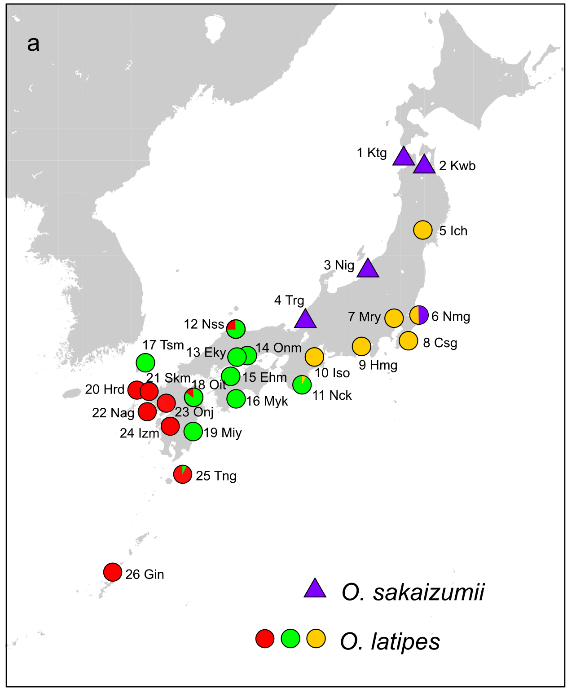

Hrd-m2

Hrd-m1

*O. latipes*, East Korea (HSOK)

b

Figure S3. (a) Geographic locations of collection sites. Solid circles and triangles on the map represent *Oryzias sakaizumii* and *Oryzias latipes*, respectively. Colors correspond to the clusters in the ADMIXTURE analysis (*K*= 4). (b) Neighbor-joining tree based on whole-genome sequences, excluding individuals with multiple ancestry in the ADMIXTURE analysis (*K* = 4). Excluded sites and individuals (Tng, Nss, Oit, Nck, and Nmg) were represented site names with underlined in the map.

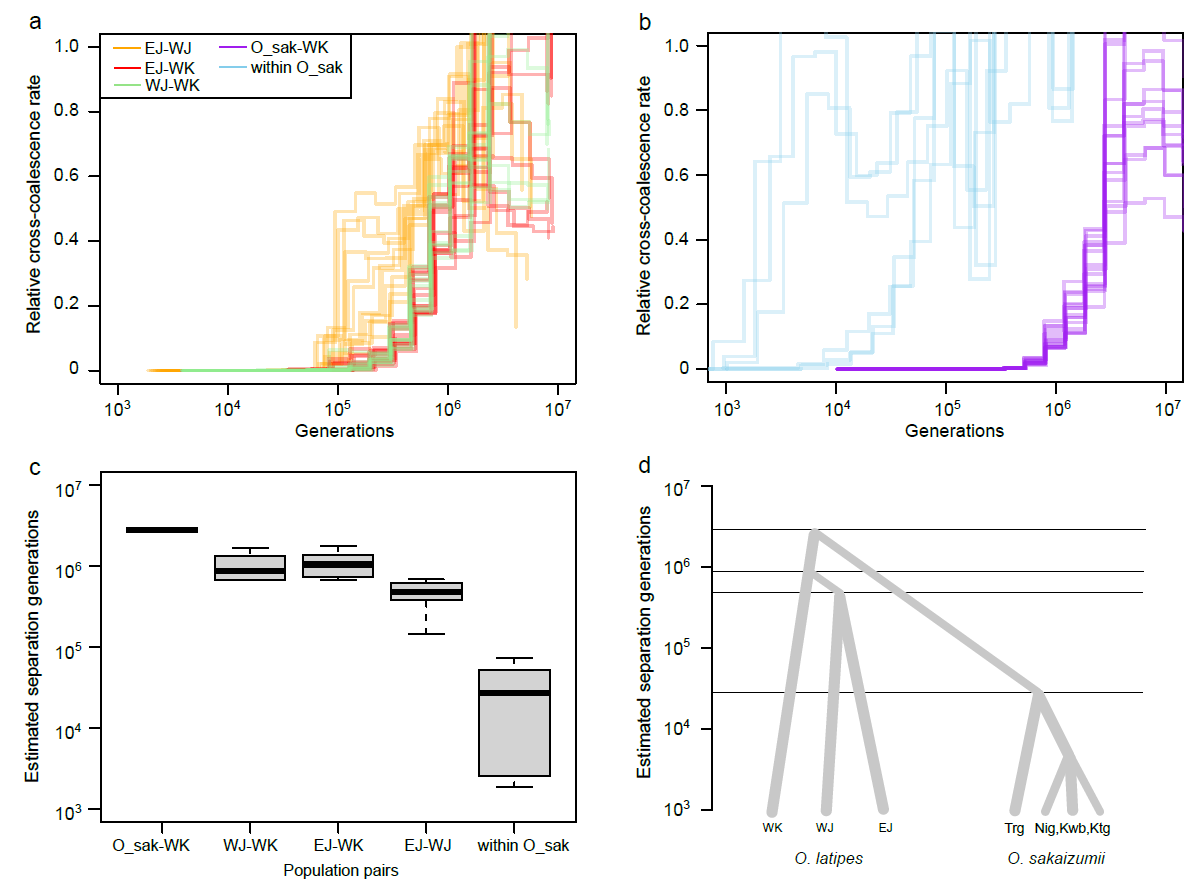

Figure S4. (a, b) Relative cross-coalescence rates between four populations (a), and between collection sites in *O. sakaizumii* (b), calculated by MSMC2 based on four individuals (eight haplotypes). (c, d) Average divergence times in generations between populations (c) and schematic representation of population divergence (d).

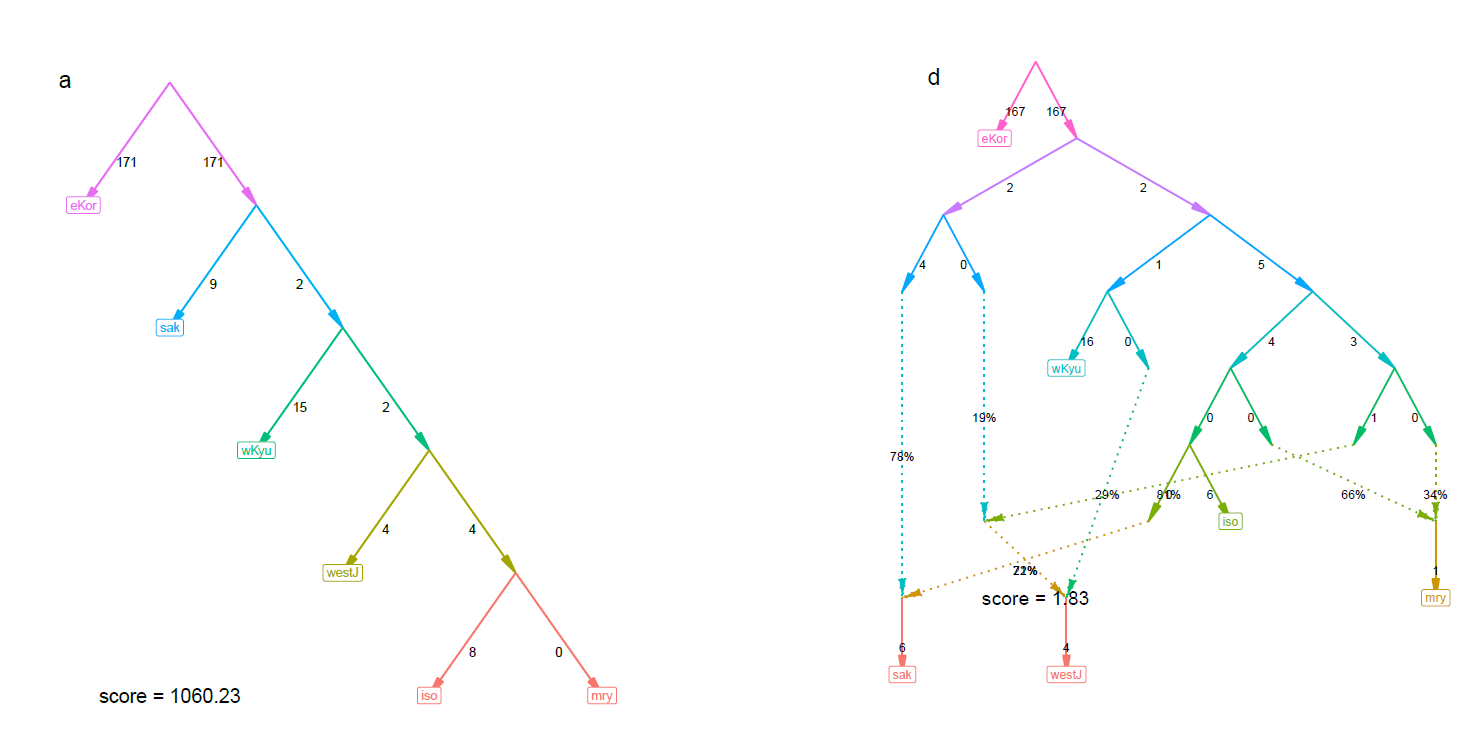

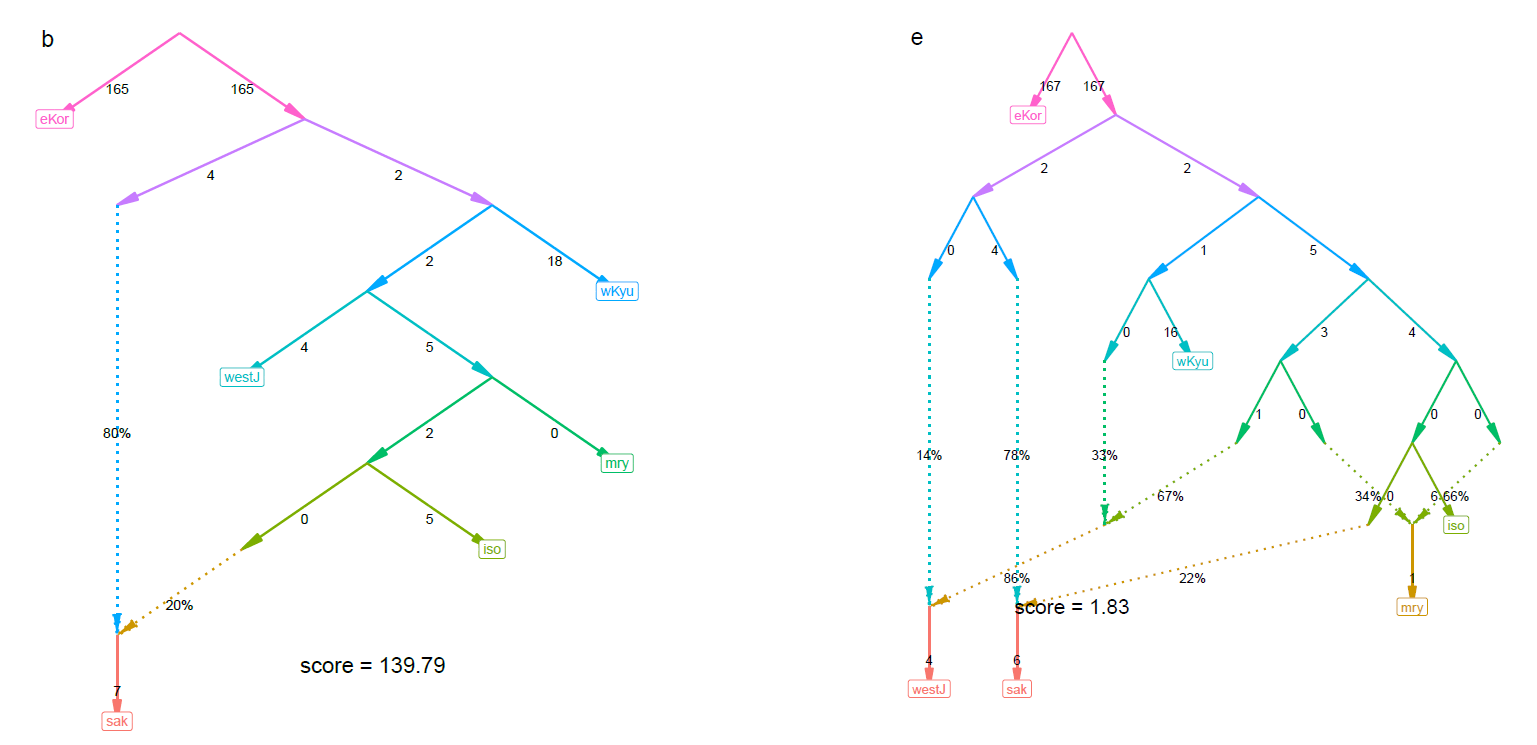

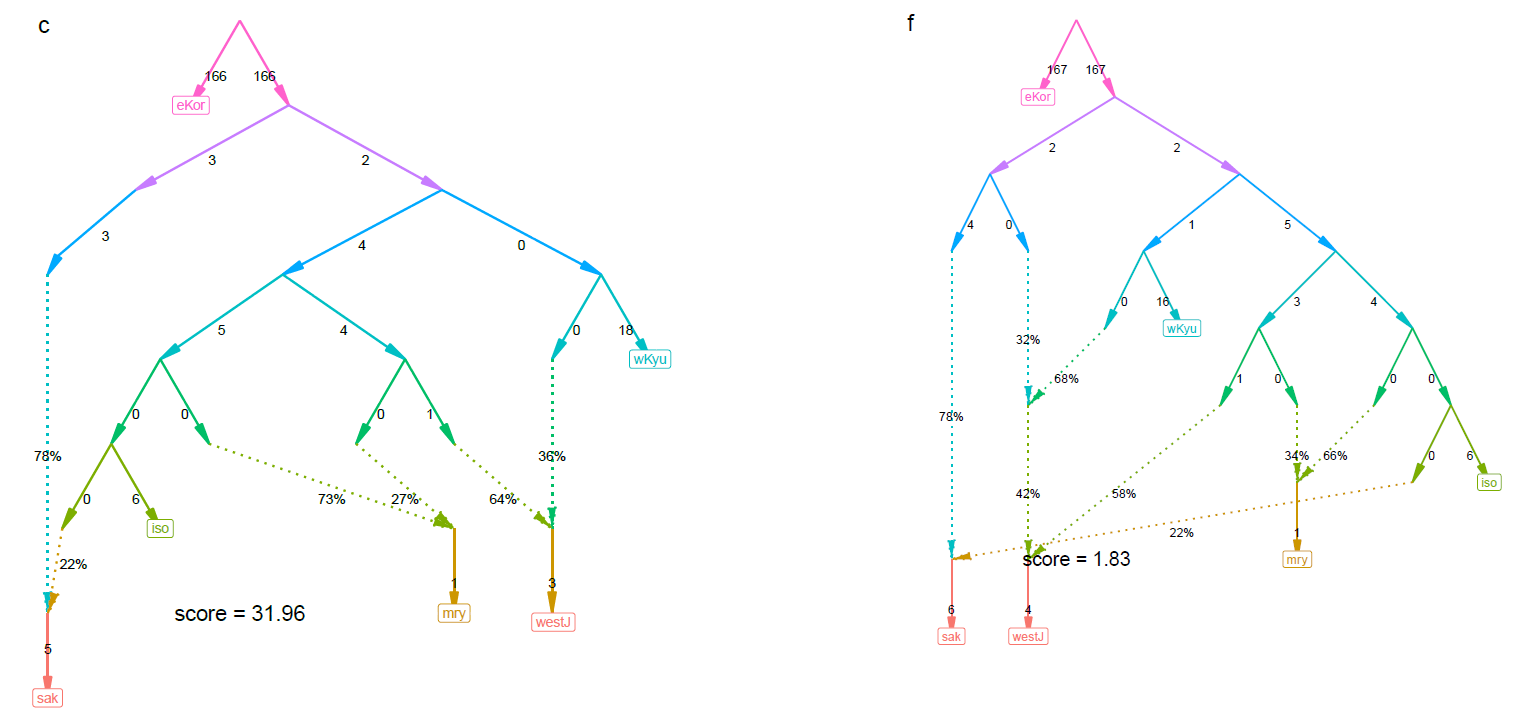

Figure S5. (a–f) Admixture graphs that fit the data. Branch lengths (*F_2_* drift distances) are shown as solid lines, and admixture events are shown as dotted lines with mixture proportions indicated. (e) Introgressive regions identified by a genomic scan of *F_d_* statistics in non-overlapping 50-SNV windows.

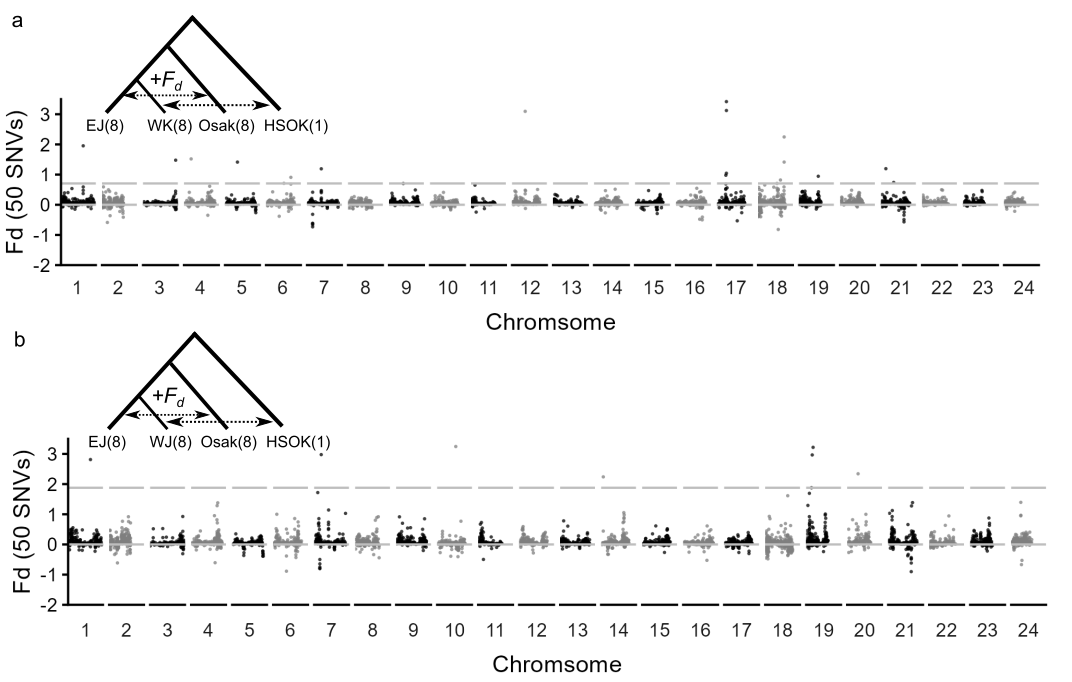

Figure S6. Signature of introgressive regions represented with positive *F_d_* values in the combination of (a) EJ, WK, Osak, HSOK and (b) EJ, WJ, Osak, HSOK. *Oryzias sakaizumii*: Osak; *Oryzias latipes*: Western Kyushu (WK), Western Japan (WJ), and Eastern Japan (EJ).

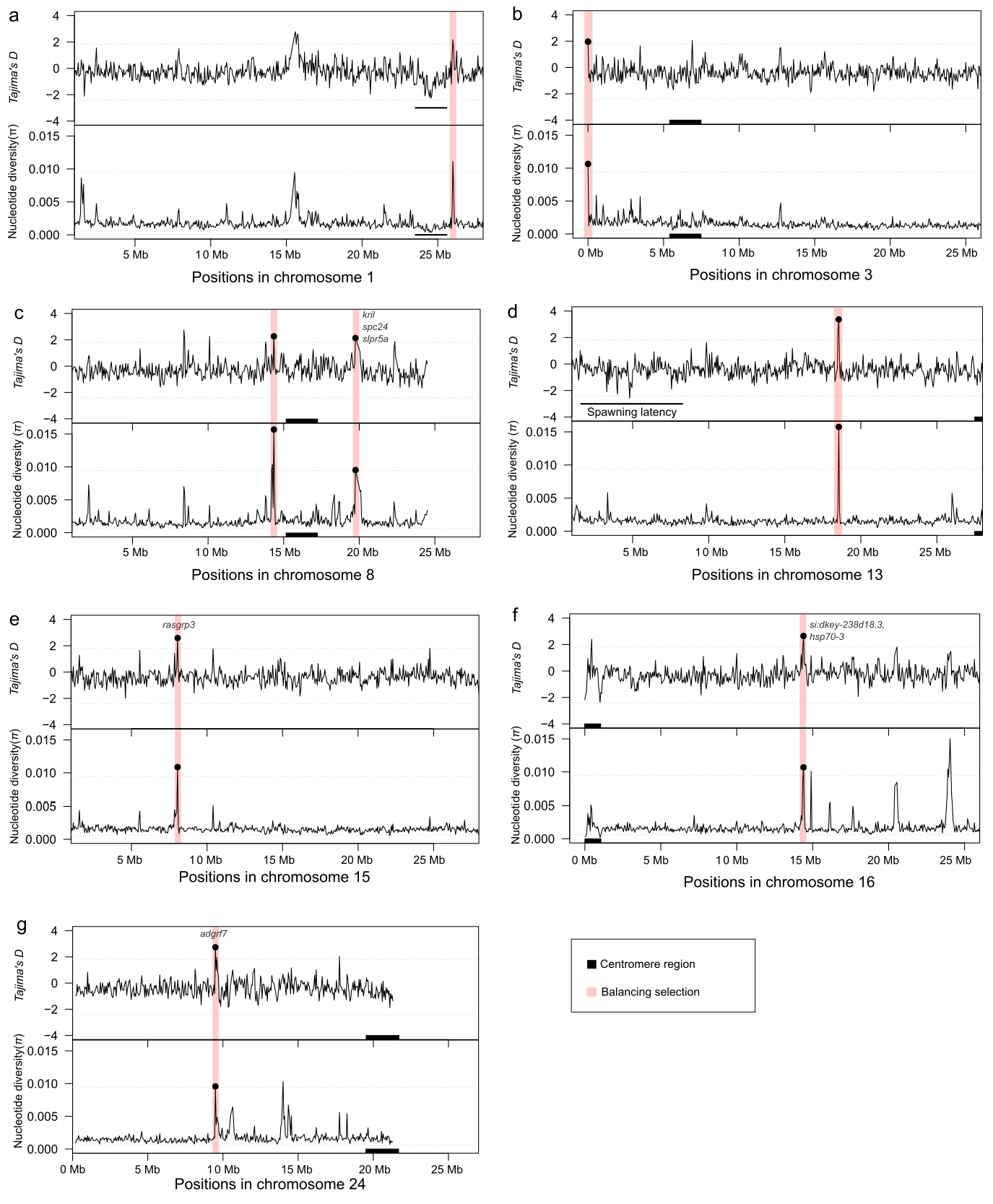

Figure. S7, Nucleotide diversity and *Tajima’s D* on chromosomes with signature of selection. Centromere regions were inferred by the synteny block positions between *Oryzias latipes* (HdrR strain, ASM223467v1, Ansai et al. 2023, Table S4) and *O. sakaizumii* (HNI strain, ASM223471v1) using ntSynt. Gene positions were based on the reference sequence of *O. sakaizumii*, HNI strain in Ensembl release 109.

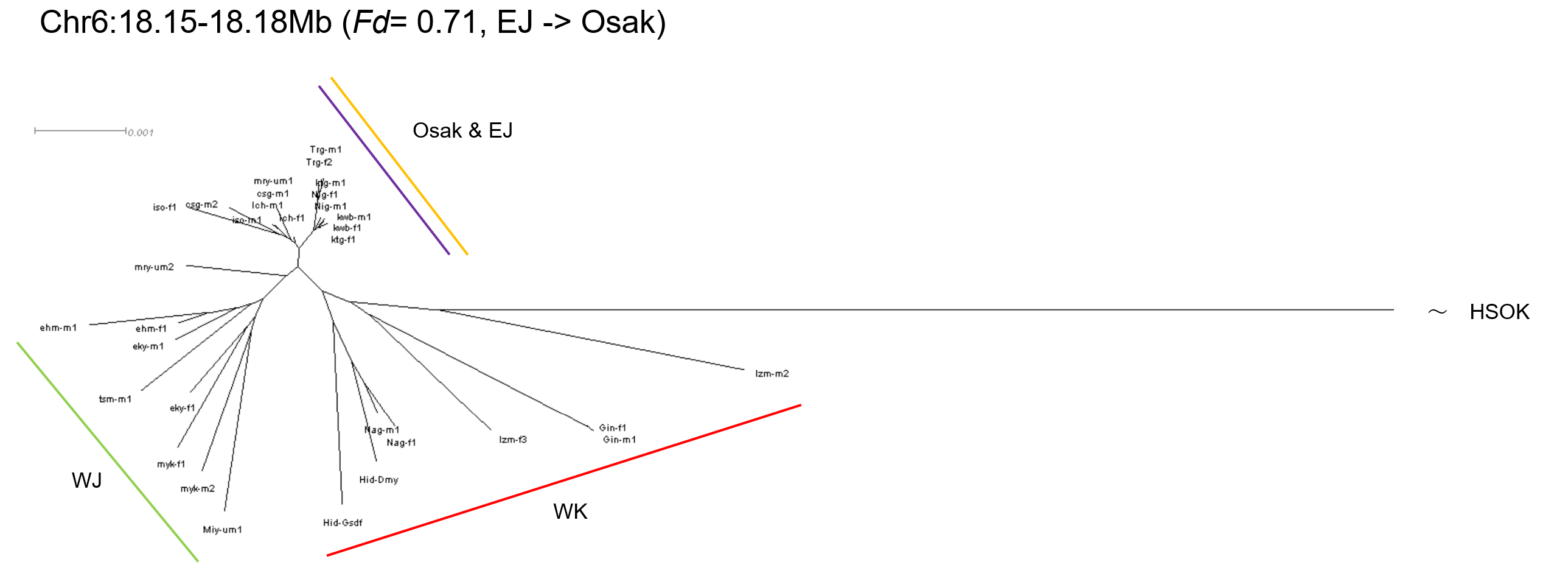

Figure. S8. Neighbor joining tree of introgressed locus on chromosome 6. *Oryzias sakaizumii*: Osak; *Oryzias latipes*: Western Kyushu (WK), Western Japan (WJ), and Eastern Japan (EJ)
